# Shotgun metagenomics reveals an enrichment of potentially cross-reactive bacterial epitopes in ankylosing spondylitis patients, as well as the effects of TNFi therapy and the host’s genotype upon microbiome composition

**DOI:** 10.1101/571430

**Authors:** Jian Yin, Peter R. Sternes, Mingbang Wang, Mark Morrison, Jing Song, Ting Li, Ling Zhou, Xin Wu, Fusheng He, Jian Zhu, Matthew A. Brown, Huji Xu

## Abstract

Diverse evidence including clinical, genetic and microbiome studies support a major role of the gut microbiome in the common immune-mediated arthropathy, ankylosing spondylitis (AS). To further investigate this we performed metagenomic analysis of a case-control cohort of 250 Han-Chinese subjects. Previous reports of gut dysbiosis in AS were re-confirmed and several notable bacterial species and functional categories were differentially abundant. TNF-inhibitor (TNFi) therapy at least partially restored the perturbed microbiome observed in untreated AS cases to that of healthy controls, including several important bacterial species that have been previously associated with AS and other related diseases. Enrichment of bacterial peptides homologous to HLA-B27-presented epitopes was observed in the stools of AS patients, suggesting that either HLA-B27 fails to clear these or that they are involved in driving HLA-B27-associated immune reactions. TNFi therapy of AS patients was also associated with a reduction of potentially arthritogenic bacterial peptides, relative to untreated patients. An AS-associated SNP in *RUNX3* significantly influenced the microbiome in two independent cohorts, highlighting a host genotype (other than *HLA-B27*) potentially influencing AS via the microbiome. These findings emphasise the key role that the gut microbiome plays in driving the pathogenesis of AS.

## INTRODUCTION

Ankylosing spondylitis (AS) is a chronic inflammatory disease affecting primarily the spine and pelvis, causing pain and initially reversible stiffness, and ultimately leading to joint ankylosis due to ectopic bone formation. In a subset of patients, peripheral joints and extra articular tissues including the eye, gut and skin are also involved. Its prevalence in Asian and European descent populations ranges from 0.09% to 0.55% ^1,2^, whereas the disease is rare in most of Africa, likely due to the low frequency of the main susceptibility gene, *HLA-B27* ^3^. There is a significant unmet therapeutic need in AS, with limited evidence that current therapies prevent spinal ankylosis, no oral treatments which suppress disease activity other than corticosteroids, and no treatments which have been demonstrated to induce remission or prevent the disease.

AS has been shown in both twin and unrelated case/control studies to be highly heritable (twins >90% heritability ^4,5^, unrelated case/control common variant heritability 69% (http://www.nealelab.is/blog/2017/9/15/heritability-of-2000-traits-and-disorders-in-the-uk-biobank)). Over past decade, at least 116 susceptibility genes have been identified, contributing 29% of the overall risk of the disease ^6^. There is substantial evidence suggesting that the interaction between host genetics and gut microbiome is a key driver of the pathogenesis of AS. The high disease heritability indicates that the environmental factors involved in the disease are likely to be ubiquitous. Reactive arthritis is a spondyloarthritis sharing many clinical and genetic features with AS, and is known to be caused by bacterial infections of the gut or urinary tract; a subset of reactive arthritis patients go on to develop AS. About 60% AS patients suffered from subclinical bowel inflammation and 10% of them can be diagnosed as inflammatory bowel disease (IBD) ^7^. There is considerable overlap in the overall heritability of AS and IBD ^8^, the two diseases are often co-familial ^9^, and many shared genetic associations have been identified ^10^. A bioinformatic study showed that AS susceptibility genes specifically enriched in gut cells are also enriched in ‘response to bacterium’ GO term pathway, and that AS-associated genetic loci are found disproportionately to lie within epigenetic marks of gene activity in gut tissue and cells ^11^. Germ-free *HLA*-*B27*-transgenic rats and SKG mice are disease-free ^12,13^. Studies using sequencing-based bacterial profiling of terminal ileal biopsies showed that AS patients have a distinct microbiome ^14^, a finding that has subsequently been reproduced studying stool samples in AS patients and patients with spondyloarthritis (SpA), a broader clinical classification ^15,16^. There has also been suggestive evidence reported that the gut microbiome is associated with differences in AS disease activity ^17^. In addition, one study compared SpA patients’ stool samples before and three months after TNF-inhibitor (TNFi) treatment onset ^18^. Although modest changes were found in microbiome alpha-diversity measures after TNFi treatment, no changes in specific bacterial taxa were observed. This may have been related to power or sampling issues, noting that 15/18 patients studied met only the ASAS axial spondyloarthritis classification criteria rather than having the more specific diagnosis, AS. In summation, the above evidence supports the contention that AS status is influenced by interactions between the gut microbiome and the host immune system.

To date, the mechanisms involved in the interaction between the host immune system and intestinal microbes remain unclear. One hypothesis suggests that HLA-B27 presents specific peptides to CD8+ T cells, leading to pathogenic adaptive immune responses (the ‘arthritogenic peptide theory’). The gut microbiota produce a huge variety and number of peptides, and as such, microbial peptides intrinsic to dysbiosis may activate CD8+ T-cells In that context, Purcell and colleagues identified 7500 such peptides that bind the eight most common *HLA-B27* subtypes ^19^. Here, we present our findings from a shotgun metagenomics sequencing study undertaken with stool samples collected from in 250 Chinese individuals, to investigate evidence of dysbiosis in AS, the effect of host genetic makeup and of TNFi treatment on the gut microbiota, and to investigate evidence of immunity to HLA-B27 restricted microbial peptides in AS cases.

## MATERIALS AND METHODS

### Subject recruitment

A total of 127 unrelated Han Chinese AS cases were recruited from the Department of Rheumatology and Immunology of Shanghai Changzheng Hospital (Shanghai, China) from December 2014 to June 2017. All cases met the 1984 modified New York criteria for AS ^20^. 123 healthy controls (blood donors on no prescription medications) were recruited from Shanghai. All human studies have been approved by the Research Ethical Committee of Second Military Medical University, and all patients and controls gave informed written consent for their participation in the studies. Clinical information was recorded for all patients, including demographic information (gender, age, smoking status and BMI), disease duration, *HLA-B27* carriage, sulfasalazine and TNFi treatment information, the Bath Ankylosing Spondylitis Disease Activity Index (BASDAI) ^21^ and Bath Ankylosing Spondylitis Functional Index (BASFI) ^22^, and clinical manifestations (inflammatory back pain, uveitis, axial arthritis, peripheral arthritis, ulcerative colitis, Crohns disease, enthesitis, dactylitis and psoriasis). Dietary habits were assessed by a 52-question questionnaire to exclude subjects with special dietary habits such as an entirely plant-based or meat-based diet. Where possible, Student’s T test and Fisher’s exact test were used to identify differences in the metadata categories between cases and controls.

### DNA microarray and subject genotyping

Samples were genotyped using the Infinium CoreExome-24v1-1 Chip (Illumina, San Diego, CA, USA) according to the manufacturer’s recommendations. Bead intensity data were processed and normalised for each sample, and genotypes called within collection using GenomeStudio.

SNPs with call rate below 95% or with a Hardy-Weinberg equilibrium of P < 10^−6^ in controls were excluded. For the overlapping SNPs, pairwise missingness tests removed all SNPs with differential missingness (P < 10^−7^). After merging data sets, SNPs with call rate below 98% and samples with call rate below 98% were removed. *HLA* alleles were imputed by SNP2HLA using the Pan-Asian reference panel^23,24^ and SNPs were extracted by PLINK v.1.90 ^25^.

### Shotgun metagenome sequencing

Faecal samples were collected and stored at −80°C prior to DNA extraction. DNA was extracted using a StoolGen DNA kit (CWBiotech Co., Beijing, China). DNA concentrations were determined using a Qubit dsDNA BR assay kit (Thermo Fisher, Foster City, CA, USA). 200 – 500 bp insert size libraries were constructed using a TruSeq DNA Sample Preparation Kit (Illumina, San Diego, CA, USA) and an automated SPRI-Works system (Beckman Coulter, San Jose, CA, USA),

Quality Control (QC) of each library was carried out using an Agilent 2100 Bioanalyzer (Agilent Technologies, Santa Clara, CA, USA), Qubit dsDNA BR assay kit (Thermo Fisher, Foster City, CA, USA) and a KAPA qPCR MasterMix plus Primer Premix kit (Kapa Biosystems, Woburn, MA, USA) according to the manufacturer’s instructions. Libraries that passed QC (>3 ng/μL) were sequenced using an Illumina HiSeq sequencer (Illumina, San Diego, CA, USA) with the paired-end 150-bp sequencing model based on >5G raw data output per sample.

Manual inspection and QC of sequencing reads was conducted using FastQC v10.1 ^26^. Paired-end reads were joined using PEAR v0.9.10 ^27^ and adapters were trimmed using Trimmomatic v0.36 ^28^. Contaminant sequences, such as those mapping to human or PhiX genomes, were filtered using Bowtie2 v2.3.4 ^29^ and the remaining reads were counted and subsampled to an equal sequencing depth of 3,520,000 sequencing reads per sample using SeqTK v1.0 ^30^. MetaPhlAn2 v2.6.0 ^31^ was used for taxonomic classification, PanPhlAn v1.2.2 ^32^ was used for strain-level profiling utilising pre-computed pan-genome references where possible, and HUMAnN2 v0.11.1 ^33^ was used for functional mapping to KEGG Orthogroups (KO) and MetaCyc pathways and utilising a UniRef90 database.

For prediction of bacterial peptides homologous to previously reported HLA-B27-presented epitopes, bacterial-derived sequencing reads were BLASTXed against a local, BLAST-formatted ^34^, version of the immune epitope database (IEDB) v3.0 (downloaded August 2016) ^35,36^. BLAST best-hits with an E-value < 0.1 were retained, annotated according to a published study ^37^ and then counted. The peptides annotated as HLA-B27-presented were compared between AS patients and healthy controls using Fisher’s exact test.

### Statistical Analysis

Abundance tables were arcsine square root transformed prior to analysis. Multidimensional data visualisation was conducted using a sparse partial least squares discriminant analysis (sPLSDA) as implemented in R as part of the MixOmics package v6.3.1 ^38^, at the species level using Bray-Curtis distance matrices. Receiver operating characteristic curve was calculated from sPLSDA using the MixOmics package v6.3.1. Controlling for covariates (such as gender, BMI, age and smoking status) where appropriate, multivariate association of the bacterial species composition with metadata of interest was conducted using a PERMANOVA test as part of vegan v2.4-5 ^39^. The alpha diversity of bacterial species was calculated using the rarefy function, as implemented in vegan v2.4-5. Univariate association of bacterial species and functional pathways/groups were tested for significance using MaAsLin v0.0.5 ^40^ and Wilcoxon rank-sum tests as implemented in R ^41^. Only results which were significant in both tests were reported in the main text. For measurement of microbial epitope richness, the Shannon, Simpson and Chao diversity indices were measured (vegan v2.4-5) and group differences were evaluated using Wilcoxon rank-sum tests.Genetic-relatedness dendrograms for strain-level results from PanPhlAn were calculated using Jaccard distance matrices and hierarchical clustering as implemented in R v.3.5.2.

## RESULTS

### Gut dysbiosis in ankylosing spondylitis

We initially sought to confirm previous reports of dysbiosis in AS cases. The case and control cohorts were divided into discovery and validation cohorts prior to analysis; the discovery cohort consisted of 97 AS cases and 93 healthy controls with age-matched demographics, and the remaining 60 subjects comprised the validation cohort (30 AS cases and 30 healthy controls) (Supplementary Table 1). With the exception of a difference in the mean age in the validation cohorts in which the controls were younger on average than cases, no differences were observed between cases and controls in either the discovery or validation cohorts. PERMANOVA and sPLSDA multivariate analysis revealed significant differentiation between the microbial composition of AS cases and healthy controls for both the discovery (P = 0.019) and validation (P = 0.0006) cohorts (Figure 1A), consistent with previous reports. Receiver-operator curve analysis showed high discrimination between cases and controls using microbiome data alone (AUC=0.87 in combined discovery and validation cohorts) (Supplementary Figure 1).

**Figure 1:**
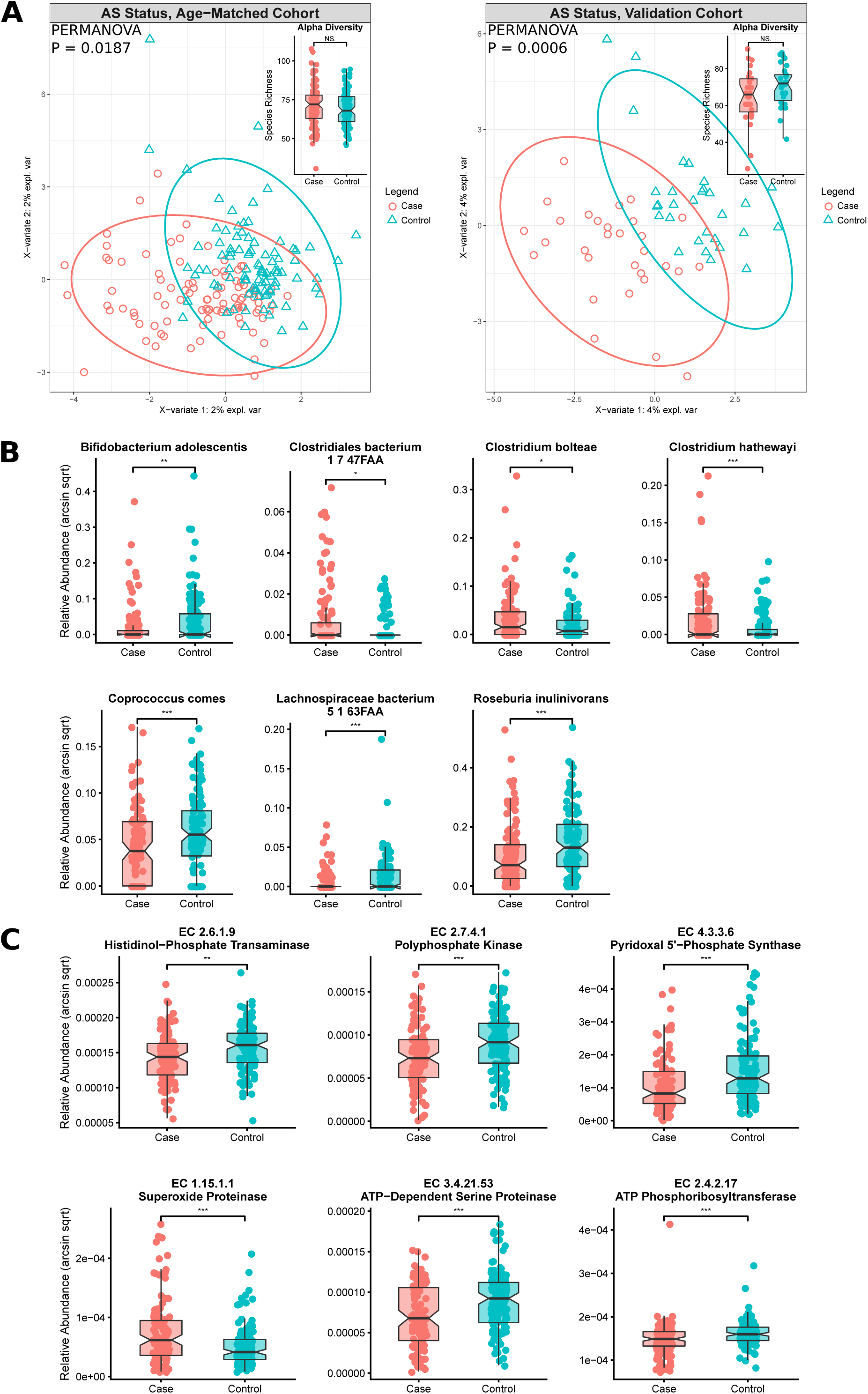
Taxonomic and functional dysbiosis observed in AS cases relative to healthy controls. **A.** Alpha and beta diversity analysis. sPLSDA and PERMANOVA revealed community-level differences in taxonomic composition. **B.** Commonly-differentiated bacterial species from the discovery and validation cohorts. **C.** Commonly-differentiated KEGG Orthogroups from the discovery and validation cohorts. Bacterial species and KEGG Orthogroups exhibiting significant results according to multivariate linear modelling and Wilcoxon rank-sum tests are shown.

Seven bacterial species were identified to be differentially abundant (P < 0.05) (i.e. were ‘indicator species’) between AS cases and healthy controls, in both the discovery and validation cohorts (Figure 1B). *Clostridiales bacterium 1 7 47FAA, Clostridium bolteae* and *Clostridium hatheway* were found to be enriched in AS cases, whilst *Bifidobacterium adolescentis, Coprococcus comes, Lachnospiraceae bacterium 5 1 63FAA* and *Roseburia inulinivorans* were depleted. Several other differentially abundant species of interest were identified in either the discovery or validation cohort, notably *Prevotella copri, Dialister invisus and Faecalibacterium prausnitzii.* A full list of differentially abundant taxa in either cohort is available in Supplementary Table 2.

Six KEGG Orthogroups were also found to be differentially abundant (P < 0.05) in both cohorts (Figure 1C), however there were no MetaCyc metabolic pathways which were differentially abundant in both cohorts. The commonly-differentiated KEGG Orthogroups were EC 2.6.1.9: histidinol-phosphate transaminase, EC 2.7.4.1: polyphosphate kinase, EC 4.3.3.6: pyridoxal 5’-phosphate synthase, EC 1.15.1.1: superoxide proteinase, EC 3.4.21.53: ATP-dependent serine phosphatase, and EC 2.4.2.17: ATP phosphoribosyltransferase. Full lists of the differentially abundant KEGG Orthogroups and MetaCyc metabolic pathways are available in Supplementary Tables 3 and 4, respectively.

Linear regression was used to investigate the correlation between the indicator species and the commonly-differentiated KEGG Orthogroups. All indicator species, except for *Lachnospiraceae bacterium 5 1 63FAA*, were significantly associated (P < 0.05) with the KEGG Orthogroups, however the degree of variation explained by these species was typically low with R^2^ values ranging from 0.0008 to 0.13 (0.043 on average) (Supplementary Table 5).

Strain-level profiling of the dysbiotic microbes identified in Figure 1B uncovered no discernible differences in strain composition between AS cases and healthy controls, with identical strains often being observed in both case and control subjects. (Supplementary Figure 2). This suggests that gut dysbiosis may primarily be a result of differential abundance at the species level and that functional or metabolic differences in the microbiome occur from common genetic elements amongst the strain population, as evidenced by KEGG Orthogroups being detectable in the majority of samples in Figure 1C.

### Effect of TNFi therapy upon the microbiome

TNFi treatment is highly effective in AS, and it is feasible that at least some of its benefits occur through effects on the gut microbiome. To test this hypothesis, the discovery and validation cohorts were combined into the following categories: healthy controls (n = 123), AS cases treated with TNFi (either etanercept or infliximab, n = 67), and AS cases who have not received TNFi treatment (n = 60). No statistically significant effect of sulfasalazine treatment was observed (P=0.76, Supplementary Figure 3). Multivariate comparison of TNFi untreated and treated cases revealed an effect of TNFi treatment upon the overall composition of the microbiome (P = 0.022) (Figure 2A). Untreated cases were significantly different to healthy controls (P = 0.0002), whereas treated cases were not significantly different to healthy controls (P = 0.069) indicating that treatment has helped restore the perturbed composition of the microbiome.

**Figure 2:**
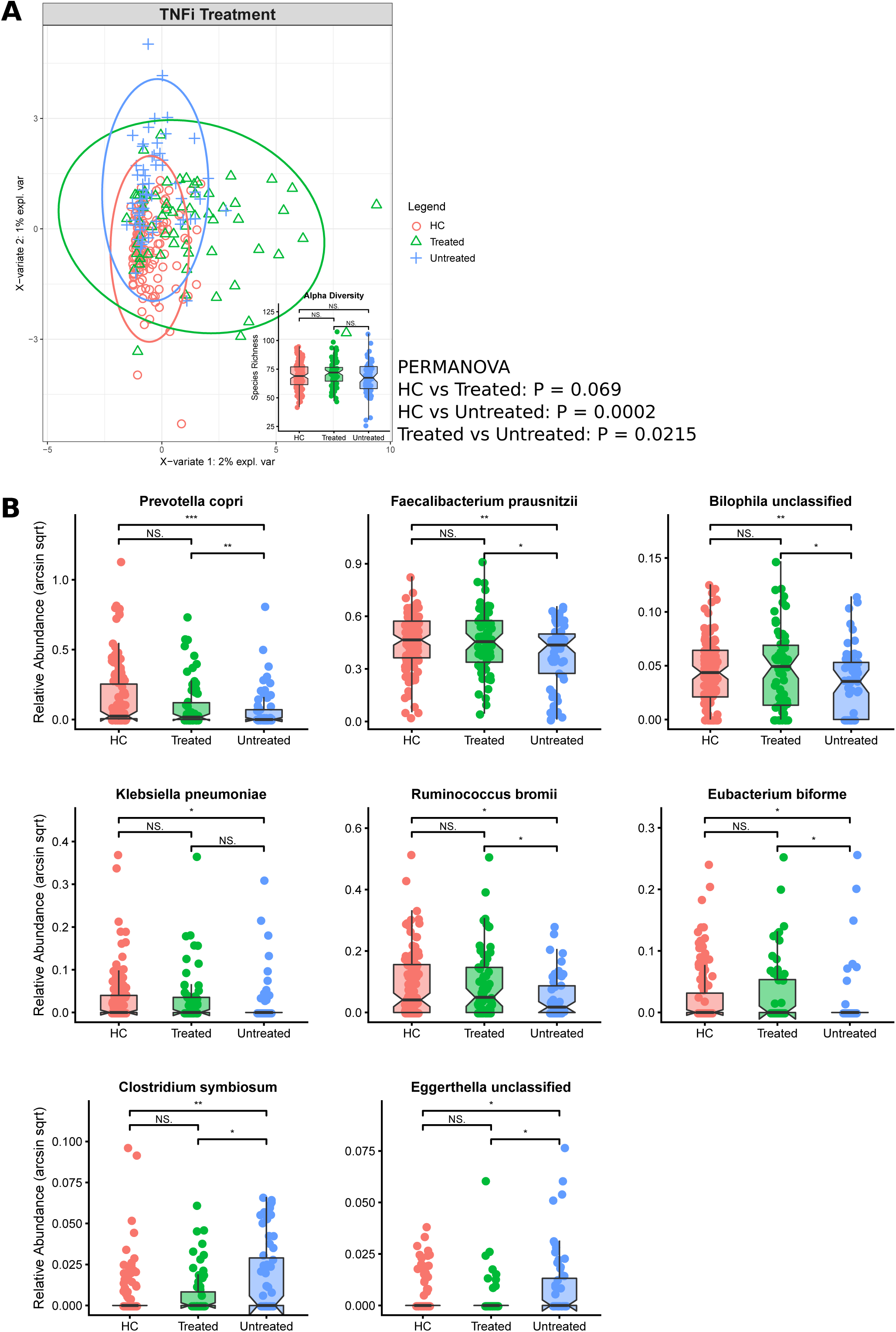
Effect of TNFi therapy upon the microbiome. **A.** Alpha and beta diversity analysis. sPLSDA and PERMANOVA revealed community-level differences in taxonomic composition. **B.** Bacterial species modulated by the effects of TNFi treatment. Bacterial species exhibiting significant results according to multivariate linear modelling and Wilcoxon rank-sum tests are shown.

To identify the key species modulated by the effects of TNFi therapy, species which were both (a) perturbed in untreated AS cases relative to healthy controls, and (b) differently abundant in treated cases compared to untreated cases, were first identified (Figure 2B and Supplementary Table 6). Six of the eight identified species exhibited significant depletion in untreated AS cases, however TNFi treatment appeared to restore the abundance of these species to levels indistinguishable from healthy controls. These species were: *Prevotella copri, Faecalibacterium prausnitzii, Bilophila unclassified, Klebsiella pneumoniae, Ruminococcus bromii* and *Eubacterium biforme*. The remaining two species (*Clostridium symbiosum* and *Eggerthella unclassified*) were enriched in untreated AS and their abundance was no longer different to healthy controls in treated cases. The findings in relation to *Prevotella copri* and *Klebsiella pneumoniae* were of particular interest given their previous association with rheumatoid arthritis (RA) and AS, respectively, as was the highly abundant (approximately 20% of total bacterial DNA, on average) *Faecalibacterium prausnitzii* for its notable depletion in several autoimmune diseases ^42^. TNFi therapy appeared to partially normalise the dysbiotic bacterial species and KEGG Orthogroups observed in AS cases relative to healthy controls shown in Figures 1B and 1C, however no statistically significant differences between treated and untreated cases were observed, potentially due to sample size constraints (Supplementary Figure 4).

The above approach was also used to identify metabolic pathways modulated by TNFi therapy. 20 MetaCyc metabolic pathways were identified in total and the perturbed abundance observed in untreated AS cases was restored to healthy control levels in 17 of these. In broad terms, these pathways primarily related to amino acid biosynthesis (notably branched-chain and aromatic amino acid biosynthesis), carbohydrate metabolism (notably starch degradation), nucleotide biosynthesis, metabolite biosynthesis and cell structure. Specific details of the 20 differentially abundant MetaCyc pathways are available in Supplementary Table 7.

Linear regression was used to investigate the association of between the modulated species and modulated pathways (Supplementary Table 8). Except for PWY-6545: Pyrimidine biosynthesis which was not associated with any individual identified species, all the pathways were significantly associated with at least two of the identified species. Similarly, all the species were significantly associated with multiple pathways, however the abundances of *Bilophila unclassified* and *Klebsiella pneumonieae* were inversely correlated with pathway abundance. An increase in *Klebsiella pneumonieae* was associated with a decrease in the abundance of PWY-6737: starch degradation (P = 0.014; R^2^ = −0.0426). The observed decrease in the starch degradation pathway for untreated AS cases is primarily attributed to a depletion of *Faecalibacterium prausnitzii* (P = 2.38×10^−24^; R^2^ = 0.3123). *Faecalibacterium prausnitzii* also exhibited strong associations with other metabolic pathways.

Strain-level profiling of the bacterial species outlined in Figure 2B also revealed no discernible differences in strain composition between healthy controls, treated cases and untreated cases, indicating the TNFi therapy affected the relative abundance of each species, not necessarily the underlying strain composition (Supplementary Figure 5).

### Effect of host genotype upon the microbiome

Genome-wide association studies (GWAS) have identified many genetic loci which are associated with AS. Emerging evidence indicates that alleles such as *HLA-B27* may influence the disease through its effect upon the gut microbiome ^43,44^. To investigate whether additional loci may affect the gut microbiome and potentially influence disease, we performed PERMANOVA analysis upon loci known to be associated with AS.

Considering non-MHC loci, an association was noted for rs11249215, a SNP in the promoter of runt-related transcription factor 3 (*RUNX3*) gene known to be associated with AS ^45,46^. This variant was associated with a shift in the microbiome of both AS cases and healthy controls (combined P = 0.0097). Furthermore, sPLSDA revealed that the degree alteration appears dependent on whether the host carried a heterozygous or homozygous genotype (Figure 3A), with the homozygous genotype resulting in a more substantial shift. As further confirmation, we analysed a recently published 16S metabarcoding dataset ^47^ of 107 healthy control subjects which were sampled from six different body sites. This analysis re-confirmed discrimination of the microbiomes based on genotype (PERMANOVA; P = 0.0001) (Figure 3B). The *RUNX3* SNP had no observable effect upon the dysbiotic bacterial species and KEGG Orthogroups outlined in Figures 1B and 1C, however its effects upon species richness and community composition (Figure 3A) suggest that further research is required to confirm a role in AS pathogenesis via effects upon the microbiome. The effect of *HLA-B27* upon the microbiome of the current cohort was unable to be investigated due to high prevalence of this genotype in AS cases and low prevalence in healthy controls, thus the effect of *HLA-B27* was unable to be discerned from the effect of AS itself.

**Figure 3:**
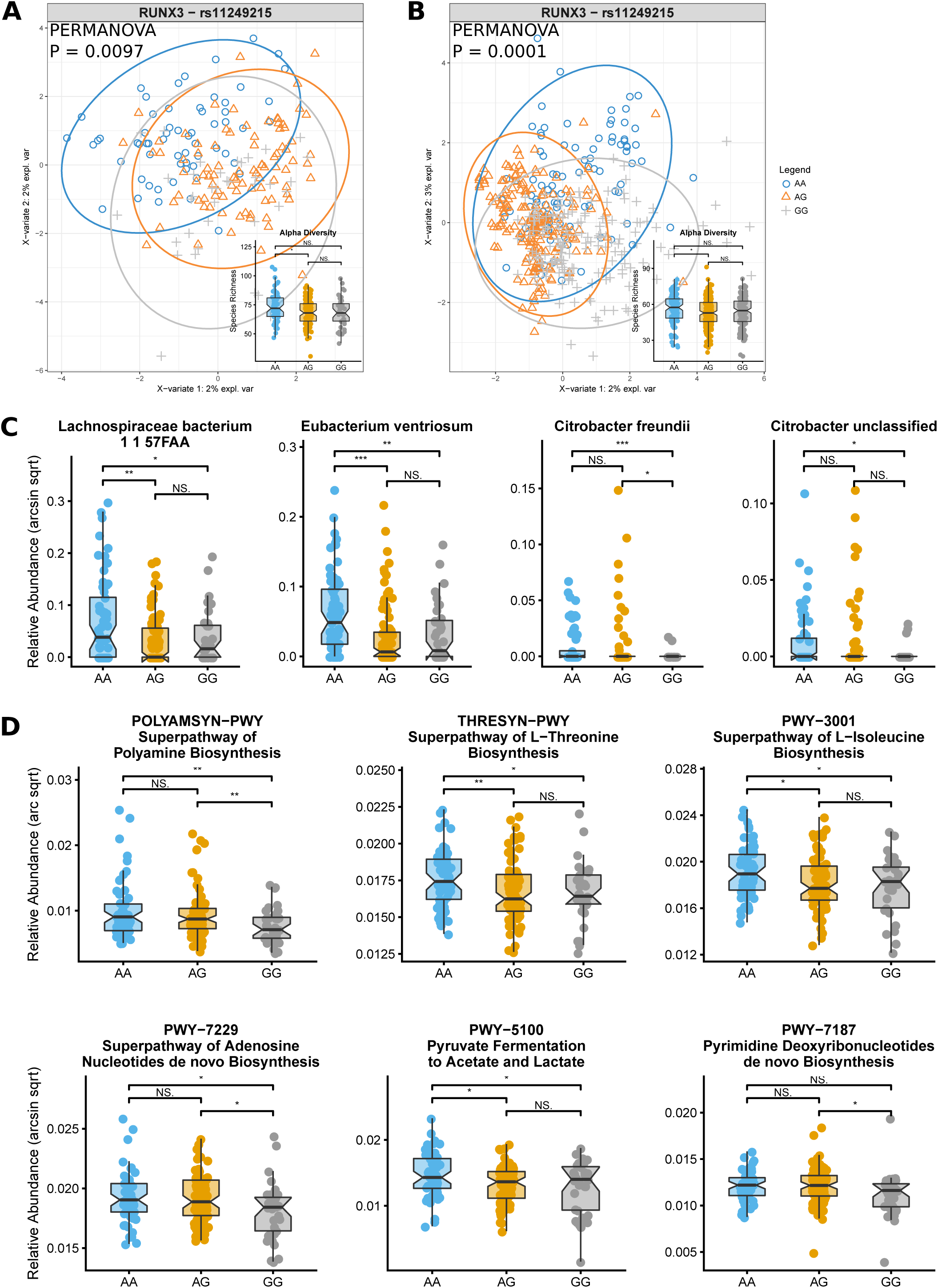
Eect of RUNX3 variants upon the microbiome. **A.** sPLSDA, alpha diversity and PERMANOVA community-level taxonomic analysis of the current study. **B.** sPSLDA, alpha diversity and PERMANOVA community-level taxonomic analysis of a recent 16S metabarcode study of healthy individuals. **C.** Modulated bacterial species according to significant results from multivariate linear modelling and Wilcoxon rank-sum testing. **D.** Modulated MetaCyc metabolic pathways, according to significant results from multivariate linear modelling and Wilcoxon rank-sum testing.

To highlight the key bacterial species affected by rs11249215, species associated with the heterozygous (AG) and homozygous (GG) genotypes, or solely the homozygous (GG) genotype were identified (Figure 3C). Several bacterial species showed differential abundance for the AG genotype, but not the GG genotype, potentially due to sample size constraints (the GG genotype was present in 37 of the 188 genotyped subjects), and thus were excluded from further analysis. Four key species identified as depleted in the AG/GG genotypes were: *Lachnospiraceae bacterium 1 1 57FAA, Eubacterium ventriosum, Citrobacter freundii* and *Citrobacter unclassified* (Supplementary Table 9).

Six MetaCyc metabolic pathways were found to be differentially abundant when comparing *RUNX3* rs11249215 genotypes (Figure 3D). Similar to the differences found for TNFi therapy, the differential pathways were primarily associated with amino acid and nucleotide biosynthesis, however notable differences in polyamine biosynthesis and pyruvate fermentation (to acetate and lactate) were also observed. Of interest is the polyamine biosynthesis pathway for the role of polyamines in enhancing the integrity of the intestinal epithelial cell barrier, and the adenosine biosynthesis pathway for the anti-inflammatory and immunosuppressant effect of adenosine (Supplementary Table 10) ^48,49^.

Linear regression of these species with the metabolic pathways revealed relatively marginal associations, with the two *Citrobacter* species exhibiting no association with any metabolic pathway (Supplementary Table 11). 16S rRNA gene metabarcoding analysis of the predominately Caucasian cohort sampled from various body sites (Figure 3B) revealed a different set of taxa and metabolic pathways potentially influenced by the *RUNX3* SNP (Supplementary Tables 12 and 13). The minimal overlap with the current shotgun metagenomic study is potentially indicative of the differences between the metagenomic approaches and/or differences in studied cohorts (i.e. geographic location, diet, ethnicity…etc).

Similar to the strain-level results for AS status and TNFi therapy, no observable bias in the underlying strain population was observed, indicating that *RUNX3* variants likely affect the relative abundance of species, not necessarily strain composition (Supplementary Figure 6^47,50^). Comparatively fewer species were associated with the *RUNX3* SNP in comparison to the number of species associated with AS status and TNFi treatment. Consistent with recent publications which have investigated the effect of the host’s genotype upon the abundance of specific taxa ^47,50^, these data provide supporting evidence that the underlying host’s genetics may have a generalised effect upon the microbiome, with a subtle effect on a higher number of taxa as opposed to a marked effect on a select few.

### Bacterial-derived HLA-B27 epitopes in AS cases and healthy controls

The main physiological function of HLA-B27 is to present peptides to CD8 lymphocytes, thereby driving cell mediated immune reactions. Differences in the presence of HLA-B27-positive epitopes in the gut microbiome in cases compared to controls, and in HLA-B27 carriers compared with HLA-B27-negative subjects, would be consistent with effects of HLA-B27 to ‘shape’ the gut microbiome, and the significance of this in regards disease pathogenesis.

To investigate the abundance of bacterial peptides homologous to HLA-B27 epitopes in AS cases and healthy controls, translated nucleotide searches were performed against IEBD v3.0, annotated according to a published study ^37^ and counted. Significant enrichment of these peptide sequences was observed in AS cases, with 24 of these enriched in both the discovery and validation cohorts (Table 1). AS cases not only exhibited enrichment of these peptides but the overall diversity of peptides was increased, with Shannon, Inverse Simpson and Chao diversity indices revealing significant differences between AS cases and healthy controls (Figure 4A).

**Table 1:**
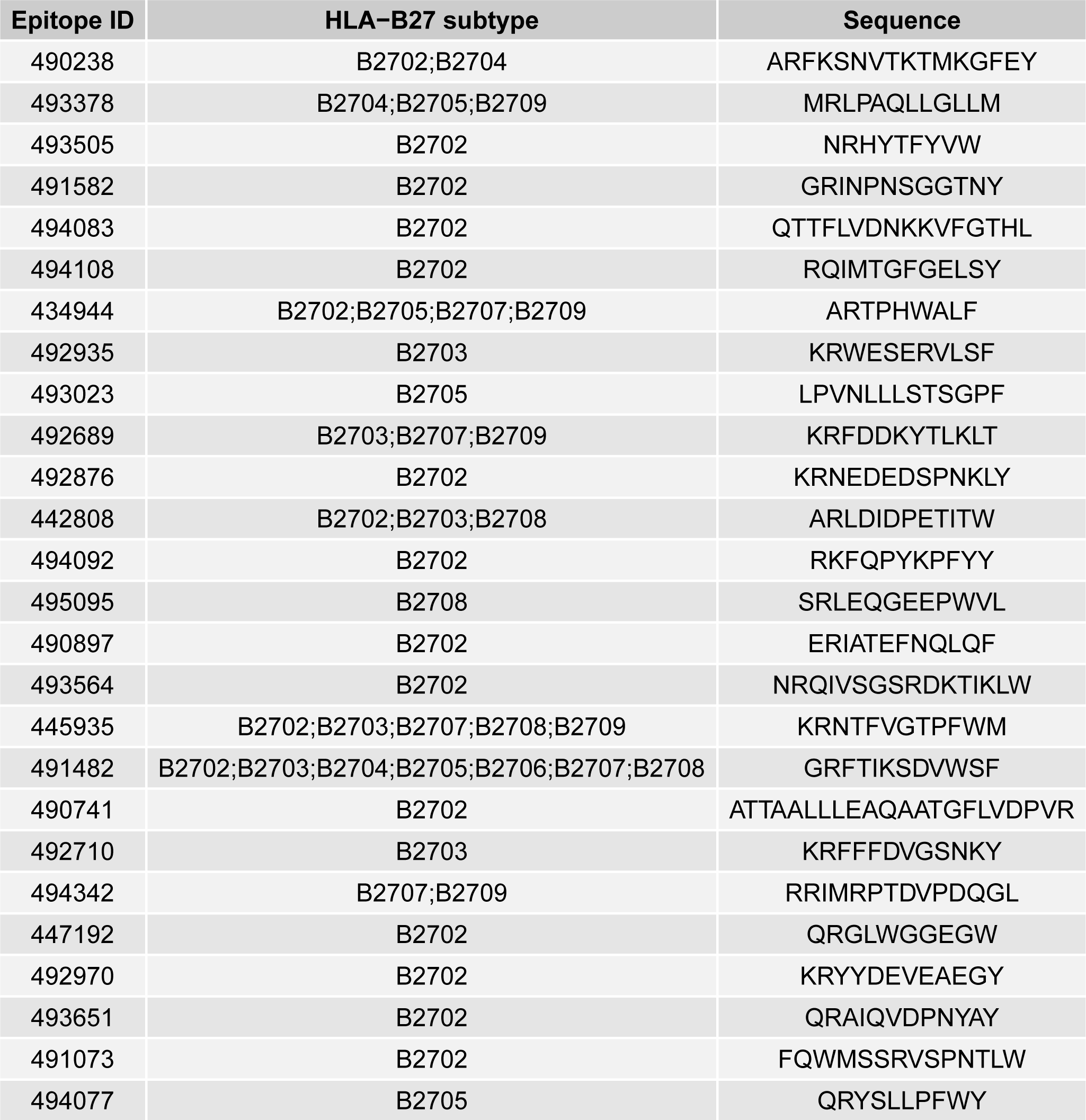
Bacterial peptides homologous to HLA-B27-presented epitopes which were commonly enriched in the discovery and validation cohorts for AS cases.

**Figure 4:**
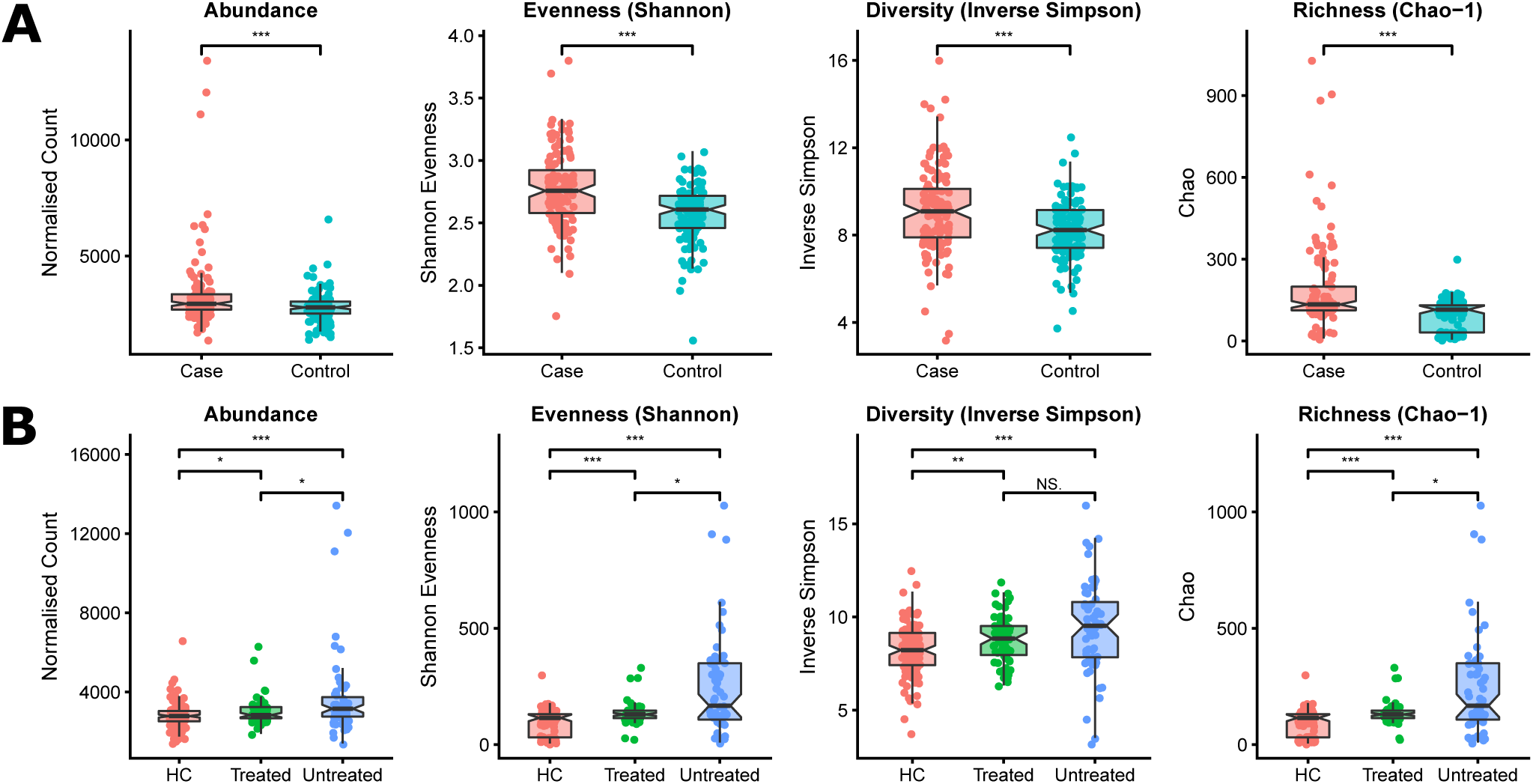
A. Enrichment, both in terms of abundance and diversity, of bacterial peptides homologous to HLA-B27 epitopes in AS cases relative to healthy controls. **B.** Differential abundance and diversity of bacterial peptides homologous to HLA-B27 epitopes in TNFi-treated and –untreated cases, and healthy controls.

TNFi treatment effects were also investigated. The overall abundance and diversity of bacterial peptides homologous to HLA-B27-presented epitopes was significantly different between the different treatment categories (Figure 4B). Untreated AS cases exhibited increased abundance and diversity of peptides. For patients who underwent TNFi therapy, a reduction in these potentially arthritogenic peptides was observed relative to untreated cases, however their levels remained marginally higher than healthy controls.

## DISCUSSION

### Gut dysbiosis in ankylosing spondylitis

This study re-confirmed the occurrence of bacterial gut dysbiosis in AS cases and identified seven bacterial species which were commonly differentiated between cases and controls in both the discovery and validation cohorts (Figure 1B). Two of these species, *Bifidobacterium adolescentis* and *Coprococcus comes*, have been noted for their depletion in Crohn’s Disease ^51^ and were also observed to be depleted in AS cases in this study. An additional two species previously reported to be associated with AS, *Prevotella copri* and *Dialister invisus*, were found to be differentially abundant in either the discovery or validation cohorts (Supplementary Table 2). In the case of *Prevotella copri*, previous studies have demonstrated enrichment in new onset RA cases yet depletion in chronic RA cases ^52^. Consistent with these findings, our study found that *Prevotella copri* was depleted in AS cases within the non-age-matched cohort, for which the demographics were heavily skewed towards older AS patients with long-standing disease (Supplementary Table 1). Previous studies in AS have shown increases in *Prevotellaceae* ^14^, or specifically with this species ^15^. As discussed below, *Prevotella copri* carriage normalised with TNFi treatment. Further studies will be required to determine if *Prevotella copri* carriage changes with disease duration, as has been reported in RA.

Carriage of *Dialister* species has been previously associated with disease activity in spondylarthrosis patients ^17^, but the carriage of *Dialister invisus* has been reported to be decreased in inflammatory bowel disease (IBD) ^53,54^. Whilst we found enrichment of *Dialister invisus* in AS cases in the discovery cohort, this was not confirmed in the validation cohort, nor has it been reported in other AS studies. Its pathogenic significance is therefore uncertain.

Of particularly interest, the notable ‘peace keeping’ microbe *Faecalibacterium prausnitzii* was also found to be depleted in AS cases in the validation cohort. This bacterium has also been consistently shown to be depleted in IBD ^54-59^, and has been previously shown to be depleted in the disease enthesitis-related arthritis, a paediatric disease-classification which includes children with ankylosing spondylitis ^60^. *Faecalibacterium prausnitzii* is known to produce butyrate and other metabolites and peptides that have diverse anti-inflammatory effects including promoting T-regulatory cell differentiation ^61^, influences on Th17 lymphocyte activation, and promotion of gut mucosal barrier function ^59,62,63^. As discussed below, TNFi treatment also led to normalisation of *Faecalibacterium prausnitzii* carriage. These findings suggest that *Faecalibacterium prausnitzii* plays a key anti-inflammatory role in AS, as it does in IBD.

The power of metagenome sequencing is to augment the widely observed phenomena collectively referred to as “ dysbiosis” beyond taxonomy-based assessment of microbiome, and provide a more highly resolved and functional characterisation of the microbiome. Here, variation in several KEGG Orthogroups remained consistent between the discovery and validation cohorts (Figure 1C). Additionally, differential abundance of some MetaCyc metabolic pathways was also observed (Supplementary Table 4), but these differences were not consistent between discovery and validation cohorts; potentially highlighting the confounding influence of the host’s age upon the metabolic composition of the gut microbiome.

Interestingly, genes encoding pyridoxal 5’-phosphate synthase, an important enzyme for the biosynthesis of vitamin B6, were less abundant amongst the microbiome of AS cases compared to healthy controls in both cohorts. Vitamin B6 plays a role in the maintenance of vitamin homeostasis in colonocytes ^64,65^. It has been found to modulate colonic inflammation and several studies have investigated the role of vitamin B6 for the treatment of inflammation in RA patients ^66-71^. Evidence from case-control studies show that RA patients have low vitamin B6 status compared to healthy controls, however intervention studies have yielded inconsistent findings, possibly due to the dose of the administered vitamin B6. The reduced potential for the microbiome of AS patients to produce pyridoxal 5’-phosphate synthase, and thus vitamin B6, may warrant further investigation into intervention strategies to mitigate inflammation in AS patients.

### Effect of TNFi therapy upon the microbiome

Previous study of RA patients before and after synthetic disease-modifying anti-rheumatic drug treatment revealed moderate differences in the gut microbiota composition, with the perturbed microbial composition being party restored following treatment ^72^. Similarly, analysis of spondylarthrosis patients before and after TNFi therapy also revealed modest differences in microbial composition yet no specific taxon was found to be modulated, likely due to sample size ^73^. Utilising a larger sample size, we confirmed that TNFi therapy was correlated with a restoration of the perturbed microbial composition, and additionally identified several notable bacterial species modulated by treatment.

We observed that TNFi therapy restored the depletion of *Faecalibacterium prausnitzii* in AS cases. Restoration of *Faecalibacterium prausnitzii* abundance was also correlated with the restored abundance of aromatic and branched-chain amino acid biosynthesis pathways. A recent ulcerative colitis study ^55^ revealed reduced dysbiosis and increased *Faecalibacterium prausnitzii* abundance in responders compared with non-responders following TNFi therapy. Furthermore, recovery of *Faecalibacterium prausnitzii* in patients with ulcerative colitis after relapse was associated with maintenance of remission 74. Another study demonstrated that treatment of infliximab completely restored *Faecalibacterium prausnitzii* concentrations from zero to 1.4 × 10^10^ bacteria/mL within few days ^52^.

Another important microbe, *Prevotella copri*, was observed to be enriched to levels closely matched to that of healthy controls following TNFi treatment. Abundance of *Prevotella copri* has previously been shown to be enriched in untreated new onset RA patients yet depleted in chronic RA cases, patients with psoriatic arthritis and healthy controls ^52^. Colonisation of SKG mice with *Prevotella copri*-dominated microbiota from RA patients exhibited increased number of Th17 cells in the large intestine ^75^. HLA-DR-presented peptides (T cell epitopes) from *Prevotella copri* were recently found to stimulate Th1 responses in 42% of new onset RA cases, with subgroups of RA patients demonstrating differential IgG or IgA immune reactivity, providing evidence that *Prevotella copri* is immune-relevant in RA pathogenesis ^76^. Additionally, the presence of the *HLA-DRB1* risk allele, which influences disease severity, in RA patients was found to be inversely correlated with *Prevotella copri* abundance ^52,77-79^. A recent study of Chinese AS patients revealed enrichment of *Prevotella copri*, as well as *Prevotella melaninogenica* and *Prevotella* sp. C561 ^15,80^. Contrasting these results, in the current study we observed depletion of *Prevotella copri* in untreated AS cases, which was restored to the healthy control levels in TNFi-treated patients. These seemingly conflicting reports of *Prevotella copri* abundance may be explained by the large degree of intraspecific genetic diversity of *Prevotella copri* strains, with strain variation adding an additional layer of complexity for predicting the function of *Prevotella copri* in the gut. The *Prevotella* genus also contains members that may be beneficial, and which do not function as pathobionts ^77-79^, with observed enrichment in healthy individuals. Taken together, our results which demonstrate a modulation of *Prevotella copri* abundance in TNFi-treated cases is a noteworthy observation, however without a stronger grasp of the strain-level genome variation within this taxon and their prevalence across our cohort, attempts to therapeutically modulate and predict the effects of *Prevotella copri* remains a significant challenge ^81^.

*Klebsiella pneumoniae* has also been suggested to play a significant role in AS pathogenesis ^82^, although this remains controversial ^83^. *Klebsiella pneumoniae* notably produces pullulanase, a starch-debranching enzyme which enables the degradation of starch into simple sugars ^84^. The apparent arthritogenic effects of dietary starch in AS are based on the concept that the growth of *Klebsiella sp.* are favoured by these diets and drive AS pathogenesis ^85,86^. Consequently, low starch diets have been promoted and are frequently followed by patients ^86^, although there is no published evidence to date as to their efficacy in positively affecting AS disease course. Here, we actually observed a depletion of this microbe in untreated cases relative to healthy controls, whereas TNFi-treated cases showed a restoration of this bacterium. Furthermore, our metagenome sequencing data showed an inverse correlation between *Klebsiella pneumoniae* relative abundance and the overall starch degradation metabolic pathway (P = 0.014; R^2^ = −0.043) (Supplementary Table 5). This pathway not only includes the pullulanase-mediated starch de-branching reaction, but also further downstream reactions including the transport and catabolism of maltodextrins. These findings do not support an association between *Klebsiella pneumoniae* and AS pathophysiology, although the role of dietary and/or resistant starches on the gut microbiota and AS warrants further investigation.

MetaCyc pathway analysis revealed depletion of the aromatic and branched-chain amino acid biosynthesis pathways in untreated AS cases, which are responsible for production of four of the nine essential amino acids in humans (leucine, isoleucine, valine and phenylalanine) (Supplementary Table 7). Vitamin B6 is an essential co-factor for branched-chain amino acid transaminase, the last step of branched-chain amino acid synthesis, and was found to be depleted in RA cases, as previously mentioned. Therefore, in the current study we not only observed a depletion of genes encoding the branched-chain amino acid biosynthesis pathway, but also for the enzyme which synthesises an important co-factor in the process.

### Enrichment of potentially arthritogenic bacterial peptides

Pathogenic bacteria have long been hypothesised as an immunological triggers of AS pathogenesis. In the current study, patients with AS not only demonstrated an enrichment of bacterial peptides matching HLA-B27 epitopes (Table 1), but the diversity of these peptides was greater overall (Figure 4A). These data provide supporting evidence for the molecular mimicry hypothesis for which bacterial-derived peptides may stimulate AS via cross-activation of autoreactive T- or B-cells, thus leading to autoimmunity. This hypothesis does not however explain the increase in HLA-B27 epitopes amongst AS cases, which could be explained by effects of non-HLA genetic factors, or AS-associated environmental factors. An alternate hypothesis is that their excess carriage is caused by a deficiency in the ability of HLA-B27 to effectively control their presence, consistent with evidence of increased bacterial migration across the gut mucosa in AS ^87^. Interestingly, the modulation of the gut microbiome caused by TNFi treatment restored the elevated abundance and diversity of peptides observed in untreated cases to levels which were more closely matched to healthy controls (Figure 4B). Further research will be required to resolve these alternate hypotheses.

### Effect of host genotype upon the microbiome

Very recently, it was demonstrated that HLA-B27 is associated with a significant shift in the microbiome in healthy individuals ^47^. Furthermore, in mouse models, MHC polymorphisms were demonstrated to contribute to an individual’s microbial composition, thus influencing health ^88^. Our investigation revealed an additional AS-associated SNP, rs11249215 in *RUNX3* ^45,46^, which was also correlated with a significant shift in bacterial composition (Figure 3A). This result was replicated in a confirmatory dataset of healthy Caucasian individuals (Figure 3B). In addition to potential roles in autoimmune diseases, variants in *RUNX3* have been associated with the intestinal inflammatory disorder celiac disease ^89^, and *RUNX3*-knockout mice spontaneously develop IBD ^90^. It is therefore tempting to hypothesise that the role of *RUNX3* in disease pathogenesis is, at least in part, caused by perturbation of the gut microbiome. Interestingly, subjects homozygous for rs11249215 exhibited a significant decrease in the abundance of the polyamine biosynthesis superpathway (Figure 3D). The intestinal tract contains high levels of polyamines which are critical for cell growth and can stimulate the production of junction proteins which are crucial for regulating paracellular permeability and reinforcing epithelial barrier function. Shifts in host and microbial polyamine metabolism may also alter the cytokine environment and induce cellular processes in both acute and chronic inflammatory settings ^49^. A potential relationship between *RUNX3*, microbial composition, intestinal polyamine levels and epithelial permeability and/or the cytokine environment warrants further investigation.

## CONCLUSION

In this study we confirm that AS is characterised by gut dysbiosis and identify key indicator species, several of which are shared with IBD. This dysbiosis is associated with functional differences in the microbiome involving known inflammation-related pathways. We demonstrate that treatment with TNFi, which is highly effective in suppressing the clinical manifestations of AS, normalises the gut microbiome, and its functional properties, in AS cases. We further demonstrate that the AS gut microbiome is enriched for bacterial peptides that have previously been shown to be presented by HLA-B27, and that this enrichment is also normalised by TNFi treatment. The impact of the host’s genotype upon microbiome composition was also highlighted, with an AS- and IBD-associated SNP in *RUNX3* correlating with a shift in microbiome composition. These findings are consistent with disease models in which AS pathogenesis is driven by interactions between a genetically primed host immune system, and the gut microbiome, and point to potential therapeutic and/or preventative approaches for the disease.

## AUTHOR CONTRIBUTIONS

Study design was performed by HX, JY and MAB. Subject recruitment and sample collection was performed by JY, JS, TL, LZ, XW and JZ. Metagenomic analysis was performed by PRS, and bacterial epitope studies by JY, FH and MW. The manuscript was prepared by PRS, MM, MAB and HX.

## ACKNOWLEDGEMENTS

HX is supported by National Science Foundation of China (Grant 81302578) and China Ministry of Science and technology (973 Program of China 2014CB541800). MAB is funded by a National Health and Medical Research Council Senior Principal Research Fellowship (APP1024879).

