## Supplementary Information for "Shotgun metagenomics reveals an enrichment of potentially cross-reactive bacterial epitopes in ankylosing spondylitis patients, as well as the effects of TNFi therapy and the host’s genotype upon microbiome composition"

**Supplementary Table 1:** Demographic and clinical characteristics of discovery and validation cohorts.

| Discovery cohort | | | |
|---|---|---|---|
| **Feature** | **Case** | **Control** | **P-value** |
| Number | 97 | 93 | - |
| % Sulfasalazine | 25.77 | 0 | - |
| % TNFi | 58.76 | 0 | - |
| Age | 35.66±11.32 | 34.46±11.65 | 0.47 |
| BMI | 22.61±2.84 | 23.18±2.55 | 0.15 |
| % Male | 80.41 | 84.95 | 0.41 |
| % B27 Pos | 96.91 | 3.63 (38 missed) | 0.00 |
| Disease Duration (yr) | 10.87±9.23 | 0 | - |
| % Smoker | 43.30 | 40.86 | 0.74 |
| BASDAI | 3.33±2.22 | 0 | - |

| Validation cohort | | | |
|---|---|---|---|
| **Feature** | **Case** | **Control** | **P-value** |
| Number | 30 | 30 | - |
| % Sulfasalazine | 26.67 | 0 | - |
| % TNFi | 33.33 | 0 | - |
| Age | 41.67±17.22 | 23.8±0.89 | 0.00 |
| BMI | 21.27±2.18 | 22.69±2.37 | 0.02 |
| % Male | 86.67 | 90 | 0.69 |
| % B27 Pos | 96.67 | 0 | 0.00 |
| Disease Duration (yr) | 13.37±11.51 | 0 | - |
| % Smoker | 30.00 | 0 | 0.00 |
| BASDAI | 3.18±2.14 | 0 | - |

**Supplementary Table 2**: Differentially abundant taxa in cases and controls in discovery and validation cohorts, identified by linear modelling.

| Feature | Coefficient | P-value |
|---|---|---|
| k__Bacteria\|p__Firmicutes\|c__Clostridia\|o__Clostridiales\|f__Peptostreptococcaceae | -0.0099 | 0.00022 |
| k__Bacteria\|p__Firmicutes\|c__Clostridia\|o__Clostridiales\|f__Peptostreptococcaceae\|g__Peptostreptococcaceae_noname | -0.0098 | 0.00024 |
| k__Bacteria\|p__Firmicutes\|c__Clostridia\|o__Clostridiales\|f__Peptostreptococcaceae\|g__Peptostreptococcaceae_noname\|s__Peptostreptococcaceae_noname_unclassified | -0.0085 | 0.0010 |
| k__Bacteria\|p__Firmicutes\|c__Clostridia\|o__Clostridiales\|f__Lachnospiraceae\|g__Roseburia\|s__Roseburia_inulinivorans | 0.040 | 0.0027 |
| k__Bacteria\|p__Firmicutes\|c__Erysipelotrichia\|o__Erysipelotrichales\|f__Erysipelotrichaceae\|g__Holdemania | -0.0071 | 0.0034 |
| k__Bacteria\|p__Firmicutes\|c__Clostridia\|o__Clostridiales\|f__Lachnospiraceae\|g__Lachnospiraceae_noname\|s__Lachnospiraceae_bacterium_5_1_63FAA | 0.0037 | 0.0063 |
| k__Bacteria\|p__Actinobacteria\|c__Actinobacteria\|o__Bifidobacteriales\|f__Bifidobacteriaceae\|g__Bifidobacterium\|s__Bifidobacterium_adolescentis | 0.012 | 0.0069 |
| k__Bacteria\|p__Firmicutes\|c__Erysipelotrichia\|o__Erysipelotrichales\|f__Erysipelotrichaceae\|g__Holdemania\|s__Holdemania_filiformis | -0.0050 | 0.013 |
| k__Bacteria\|p__Proteobacteria\|c__Betaproteobacteria\|o__Burkholderiales\|f__Sutterellaceae\|g__Parasutterella | 0.0062 | 0.014 |
| k__Bacteria\|p__Proteobacteria\|c__Betaproteobacteria\|o__Burkholderiales\|f__Sutterellaceae\|g__Parasutterella\|s__Parasutterella_excrementihominis | 0.0062 | 0.014 |
| k__Bacteria\|p__Firmicutes\|c__Clostridia\|o__Clostridiales\|f__Clostridiaceae\|g__Clostridium\|s__Clostridium_hathewayi | -0.0052 | 0.016 |
| k__Bacteria\|p__Proteobacteria\|c__Betaproteobacteria\|o__Burkholderiales\|f__Burkholderiales_noname | 0.0060 | 0.016 |
| k__Bacteria\|p__Proteobacteria\|c__Betaproteobacteria\|o__Burkholderiales\|f__Burkholderiales_noname\|g__Burkholderiales_noname | 0.0060 | 0.016 |
| k__Bacteria\|p__Proteobacteria\|c__Betaproteobacteria\|o__Burkholderiales\|f__Burkholderiales_noname\|g__Burkholderiales_noname\|s__Burkholderiales_bacterium_1_1_47 | 0.0060 | 0.016 |
| k__Bacteria\|p__Proteobacteria\|c__Gammaproteobacteria\|o__Enterobacteriales\|f__Enterobacteriaceae\|g__Enterobacter\|s__Enterobacter_cloacae | 0.013 | 0.017 |
| k__Bacteria\|p__Firmicutes\|c__Clostridia\|o__Clostridiales\|f__Clostridiales_noname | -0.0050 | 0.025 |
| k__Bacteria\|p__Firmicutes\|c__Clostridia\|o__Clostridiales\|f__Lachnospiraceae\|g__Coprococcus\|s__Coprococcus_comes | 0.013 | 0.025 |
| k__Bacteria\|p__Firmicutes\|c__Clostridia\|o__Clostridiales\|f__Clostridiales_noname\|g__Clostridiales_noname | -0.0036 | 0.030 |
| k__Bacteria\|p__Firmicutes\|c__Clostridia\|o__Clostridiales\|f__Clostridiales_noname\|g__Clostridiales_noname\|s__Clostridiales_bacterium_1_7_47FAA | -0.0036 | 0.030 |
| k__Bacteria\|p__Bacteroidetes\|c__Bacteroidia\|o__Bacteroidales\|f__Porphyromonadaceae\|g__Odoribacter\|s__Odoribacter_splanchnicus | -0.0095 | 0.030 |
| k__Bacteria\|p__Firmicutes\|c__Negativicutes\|o__Selenomonadales\|f__Veillonellaceae\|g__Megamonas\|s__Megamonas_rupellensis | -0.0023 | 0.035 |
| k__Bacteria\|p__Firmicutes\|c__Negativicutes\|o__Selenomonadales\|f__Veillonellaceae\|g__Dialister\|s__Dialister_invisus | -0.022 | 0.043 |
| k__Bacteria\|p__Firmicutes\|c__Clostridia\|o__Clostridiales\|f__Lachnospiraceae\|g__Blautia\|s__Ruminococcus_obeum | 0.0042 | 0.046 |
| k__Bacteria\|p__Firmicutes\|c__Clostridia\|o__Clostridiales\|f__Lachnospiraceae\|g__Coprococcus | 0.018 | 0.047 |
| k__Bacteria\|p__Firmicutes\|c__Clostridia\|o__Clostridiales\|f__Clostridiaceae\|g__Clostridium\|s__Clostridium_bolteae | -0.0075 | 0.050 |

| Feature | Coefficient | P-value |
|---|---|---|
| k__Bacteria\|p__Actinobacteria\|c__Actinobacteria\|o__Bifidobacteriales\|f__Bifidobacteriaceae\|g__Bifidobacterium\|s__Bifidobacterium_adolescentis | 0.055 | 0.00059 |
| k__Bacteria\|p__Bacteroidetes\|c__Bacteroidia\|o__Bacteroidales\|f__Bacteroidaceae\|g__Bacteroides\|s__Bacteroides_thetaiotaomicron | -0.044 | 0.00071 |
| k__Bacteria\|p__Firmicutes\|c__Clostridia\|o__Clostridiales\|f__Clostridiaceae\|g__Clostridium\|s__Clostridium_hathewayi | -0.0074 | 0.0040 |
| k__Bacteria\|p__Firmicutes\|c__Clostridia\|o__Clostridiales\|f__Lachnospiraceae\|g__Dorea | 0.036 | 0.0042 |
| k__Bacteria\|p__Firmicutes\|c__Clostridia\|o__Clostridiales\|f__Lachnospiraceae\|g__Lachnospiraceae_noname\|s__Lachnospiraceae_bacterium_5_1_63FAA | 0.011 | 0.0069 |
| k__Bacteria\|p__Firmicutes\|c__Clostridia\|o__Clostridiales\|f__Lachnospiraceae\|g__Dorea\|s__Dorea_longicatena | 0.021 | 0.0071 |
| k__Bacteria\|p__Firmicutes\|c__Clostridia\|o__Clostridiales\|f__Lachnospiraceae\|g__Butyrivibrio | 0.011 | 0.010 |
| k__Bacteria\|p__Firmicutes\|c__Clostridia\|o__Clostridiales\|f__Lachnospiraceae\|g__Butyrivibrio\|s__Butyrivibrio_unclassified | 0.011 | 0.010 |
| k__Bacteria\|p__Firmicutes\|c__Clostridia\|o__Clostridiales\|f__Lachnospiraceae\|g__Coprococcus\|s__Coprococcus_comes | 0.029 | 0.011 |
| k__Bacteria\|p__Firmicutes\|c__Clostridia\|o__Clostridiales\|f__Ruminococcaceae\|g__Anaerotruncus\|s__Anaerotruncus_colihominis | -0.0082 | 0.013 |
| k__Bacteria\|p__Firmicutes | 0.15 | 0.013 |
| k__Bacteria\|p__Actinobacteria\|c__Actinobacteria\|o__Actinomycetales\|f__Micrococcaceae | -0.0077 | 0.015 |
| k__Bacteria\|p__Actinobacteria\|c__Actinobacteria\|o__Actinomycetales\|f__Micrococcaceae\|g__Rothia | -0.0077 | 0.015 |
| k__Bacteria\|p__Actinobacteria\|c__Actinobacteria\|o__Actinomycetales | -0.0084 | 0.016 |
| k__Bacteria\|p__Actinobacteria\|c__Actinobacteria\|o__Actinomycetales\|f__Micrococcaceae\|g__Rothia\|s__Rothia_mucilaginosa | -0.0073 | 0.017 |
| k__Bacteria\|p__Firmicutes\|c__Clostridia\|o__Clostridiales\|f__Clostridiales_noname\|g__Clostridiales_noname | -0.0082 | 0.018 |
| k__Bacteria\|p__Firmicutes\|c__Clostridia\|o__Clostridiales\|f__Clostridiales_noname\|g__Clostridiales_noname\|s__Clostridiales_bacterium_1_7_47FAA | -0.0082 | 0.018 |
| k__Bacteria\|p__Actinobacteria\|c__Actinobacteria\|o__Bifidobacteriales\|f__Bifidobacteriaceae\|g__Bifidobacterium\|s__Bifidobacterium_pseudocatenulatum | 0.022 | 0.018 |
| k__Bacteria\|p__Bacteroidetes\|c__Bacteroidia\|o__Bacteroidales\|f__Bacteroidaceae | -0.13 | 0.019 |
| k__Bacteria\|p__Bacteroidetes\|c__Bacteroidia\|o__Bacteroidales\|f__Bacteroidaceae\|g__Bacteroides | -0.13 | 0.019 |
| k__Bacteria\|p__Firmicutes\|c__Clostridia | 0.14 | 0.020 |
| k__Bacteria\|p__Firmicutes\|c__Clostridia\|o__Clostridiales | 0.14 | 0.020 |
| k__Bacteria\|p__Firmicutes\|c__Negativicutes\|o__Selenomonadales\|f__Veillonellaceae\|g__Megasphaera\|s__Megasphaera_unclassified | 0.023 | 0.033 |
| k__Bacteria\|p__Firmicutes\|c__Clostridia\|o__Clostridiales\|f__Lachnospiraceae\|g__Dorea\|s__Dorea_formicigenerans | 0.025 | 0.035 |
| k__Bacteria\|p__Firmicutes\|c__Clostridia\|o__Clostridiales\|f__Lachnospiraceae\|g__Roseburia\|s__Roseburia_inulinivorans | 0.070 | 0.038 |
| k__Bacteria\|p__Firmicutes\|c__Clostridia\|o__Clostridiales\|f__Clostridiaceae\|g__Clostridium\|s__Clostridium_bolteae | -0.0071 | 0.038 |
| k__Bacteria\|p__Bacteroidetes\|c__Bacteroidia\|o__Bacteroidales\|f__Prevotellaceae\|g__Paraprevotella\|s__Paraprevotella_unclassified | 0.013 | 0.041 |
| k__Bacteria\|p__Bacteroidetes\|c__Bacteroidia\|o__Bacteroidales\|f__Prevotellaceae\|g__Prevotella\|s__Prevotella_copri | 0.0046 | 0.041 |

| | | |
|---|---|---|
| k__Bacteria\|p__Proteobacteria\|c__Betaproteobacteria\|o__Burkholderiales\|f__Burkholderiales_noname | 0.014 | 0.047 |
| k__Bacteria\|p__Bacteroidetes\|c__Bacteroidia\|o__Bacteroidales\|f__Prevotellaceae\|g__Paraprevotella\|s__Paraprevotella_clara | 0.012 | 0.049 |

**Supplementary Table 3:** KEGG Orthogroup comparisons in cases and controls in discovery and validation cohorts, identified by linear modelling.

| Discovery cohort | | |
|---|---|---|
| **Feature** | **Coefficient** | **P-value** |
| TRIOSEPISOMERIZATION-RXN | -0.00057 | 0.0017 |
| TRYPTOPHAN--TRNA-LIGASE-RXN | -0.00065 | 0.0025 |
| 3.1.22.4-RXN | -0.00057 | 0.0056 |
| RXN-12587 | 0.00050 | 0.0056 |
| RXN-12588 | 0.00050 | 0.0056 |
| RXN-14384 | 0.00050 | 0.0056 |
| RXN-14385 | 0.00050 | 0.0056 |
| RXN-14386 | 0.00050 | 0.0056 |
| RXN-15881 | 0.00050 | 0.0056 |
| RXN-9787 | 0.00050 | 0.0056 |
| RXN0-308 | 0.00050 | 0.0056 |
| 3.1.11.6-RXN | -0.0013 | 0.0079 |
| 3.5.1.80-RXN | -0.00065 | 0.0089 |
| NAG6PDEACET-RXN | -0.00065 | 0.0089 |
| ACETOLACTREDUCTOISOM-RXN | -0.00051 | 0.0093 |
| ACETOOHBUTREDUCTOISOM-RXN | -0.00051 | 0.0093 |
| GLURS-RXN | -0.00033 | 0.0098 |
| 3.4.21.53-RXN | 0.00081 | 0.012 |
| RXN-11322 | 0.0014 | 0.012 |
| ATPPHOSPHORIBOSYLTRANS-RXN | 0.00039 | 0.015 |
| SERINE-O-ACETTRAN-RXN | -0.00052 | 0.018 |
| HOLO-ACP-SYNTH-RXN | -0.00076 | 0.022 |
| RXN-10994 | -0.00076 | 0.022 |
| RXN-16759 | -0.00076 | 0.022 |
| 4.2.99.18-RXN | -0.00036 | 0.023 |
| RXN0-2601 | -0.00036 | 0.023 |
| 5.1.3.9-RXN | -0.00062 | 0.024 |
| NANE-RXN | -0.00062 | 0.024 |
| HOMOSERKIN-RXN | -0.00058 | 0.026 |
| ENTDB-RXN | -0.00075 | 0.029 |
| RXN-15889 | -0.00075 | 0.029 |
| HISTAMINOTRANS-RXN | 0.00045 | 0.031 |
| SUPEROX-DISMUT-RXN | -0.00080 | 0.032 |
| DEOXYRIBOSE-P-ALD-RXN | -0.00063 | 0.035 |
| POLYPHOSPHATE-KINASE-RXN | 0.00056 | 0.037 |
| SHIKIMATE-KINASE-RXN | 0.00038 | 0.047 |
| DTMPKI-RXN | -0.00054 | 0.048 |

| Validation cohort | | |
|---|---|---|
| **Feature** | **Coefficient** | **P-value** |
| 2.3.1.179-RXN | 0.0025 | 8.56E-06 |
| RXN-8391 | 0.0025 | 8.56E-06 |
| RXN1G-1015 | 0.0025 | 8.56E-06 |
| RXN1G-1212 | 0.0025 | 8.56E-06 |
| RXN1G-172 | 0.0025 | 8.56E-06 |

| | | |
|---|---|---|
| RXN1G-202 | 0.0025 | 8.56E-06 |
| RXN1G-32 | 0.0025 | 8.56E-06 |
| RXN1G-334 | 0.0025 | 8.56E-06 |
| RXN1G-374 | 0.0025 | 8.56E-06 |
| RXN1G-424 | 0.0025 | 8.56E-06 |
| RXN1G-557 | 0.0025 | 8.56E-06 |
| RXN1G-580 | 0.0025 | 8.56E-06 |
| RXN1G-582 | 0.0025 | 8.56E-06 |
| RXN1G-660 | 0.0025 | 8.56E-06 |
| RXN1G-79 | 0.0025 | 8.56E-06 |
| RXN1G-818 | 0.0025 | 8.56E-06 |
| RXN1G-883 | 0.0025 | 8.56E-06 |
| RXN1G-89 | 0.0025 | 8.56E-06 |
| RXN1G-94 | 0.0025 | 8.56E-06 |
| GLYC3PDEHYDROGBIOSYN-RXN | 0.0010 | 2.78E-05 |
| RXN-11637 | 0.0011 | 9.14E-05 |
| RXN-11135 | 0.0020 | 0.00017 |
| DNA-DIRECTED-DNA-POLYMERASE-RXN | 0.0024 | 0.00022 |
| GLYCOGEN-BRANCH-RXN | 0.0028 | 0.00024 |
| RXN-14371 | 0.0028 | 0.00024 |
| RXN-14372 | 0.0028 | 0.00024 |
| RXN-7669 | 0.0028 | 0.00024 |
| RXN-7710 | 0.0028 | 0.00024 |
| PANTOTHENATE-KIN-RXN | 0.0010 | 0.00026 |
| SUPEROX-DISMUT-RXN | -0.0026 | 0.00030 |
| 3.4.21.53-RXN | 0.0025 | 0.00037 |
| TRNA-GUANINE-N7--METHYLTRANSFERASE-RXN | 0.0011 | 0.00047 |
| HOMSUCTRAN-RXN | 0.0009 | 0.00060 |
| RXN-11322 | 0.0038 | 0.00069 |
| RXN-12458 | 0.0011 | 0.00069 |
| RXN-14517 | 0.0011 | 0.00069 |
| 2.7.1.148-RXN | 0.0011 | 0.00076 |
| N-ACETYLGLUTPREDUCT-RXN | 0.0010 | 0.00086 |
| RXN-15006 | 0.0010 | 0.00086 |
| MALONYL-COA-ACP-TRANSACYL-RXN | 0.00089 | 0.00093 |
| RXN-8850 | 0.0022 | 0.00095 |
| RXN-11638 | 0.0012 | 0.00098 |
| 3-DEHYDROQUINATE-DEHYDRATASE-RXN | 0.0011 | 0.00100 |
| POLYPHOSPHATE-KINASE-RXN | 0.0022 | 0.0011 |
| ATPSYN-RXN | 0.0025 | 0.0012 |
| DIHYDROXYISOVALDEHYDRAT-RXN | 0.0014 | 0.0013 |
| DIHYDROXYMETVALDEHYDRAT-RXN | 0.0014 | 0.0013 |
| METHIONYL-TRNA-FORMYLTRANSFERASE-RXN | 0.0009 | 0.0013 |
| AMINOCYL-TRNA-HYDROLASE-RXN | 0.0010 | 0.0014 |
| RXN-12460 | 0.0010 | 0.0014 |
| RXN-16637 | 0.0010 | 0.0014 |
| ATPPHOSPHORIBOSYLTRANS-RXN | 0.0011 | 0.0014 |
| ADENOSYLHOMOCYSTEINE-NUCLEOSIDASE-RXN | 0.0012 | 0.0015 |
| TRNA-PSEUDOURIDINE-SYNTHASE-I-RXN | 0.0018 | 0.0015 |
| ABC-27-RXN | 0.0011 | 0.0015 |

| | | |
|---|---|---|
| RXN-13997 | 0.0009 | 0.0017 |
| RXN-11633 | 0.0010 | 0.0017 |
| 6.3.5.7-RXN | 0.0017 | 0.0018 |
| TAGAKIN-RXN | 0.0020 | 0.0021 |
| HISTAMINOTRANS-RXN | 0.0013 | 0.0021 |
| PGLUCISOM-RXN | 0.00080 | 0.0022 |
| RXN-13720 | 0.00080 | 0.0022 |
| RXN-6182 | 0.00080 | 0.0022 |
| ADENYL-KIN-RXN | 0.0011 | 0.0023 |
| NICONUCADENYLYLTRAN-RXN | 0.00093 | 0.0026 |
| RXN-11592 | 0.0012 | 0.0027 |
| 1.5.1.20-RXN | 0.00088 | 0.0028 |
| 3.1.26.3-RXN | 0.00051 | 0.0031 |
| ORNCARBAMTRANSFER-RXN | 0.0017 | 0.0032 |
| RXN-13482 | 0.0017 | 0.0032 |
| DIHYDRODIPICSYN-RXN | 0.0012 | 0.0032 |
| L-LACTATE-DEHYDROGENASE-RXN | 0.0019 | 0.0033 |
| DMBPPRIBOSYLTRANS-RXN | 0.0010 | 0.0034 |
| 1.8.1.4-RXN | -0.0021 | 0.0035 |
| RXN-7716 | -0.0021 | 0.0035 |
| RXN-7719 | -0.0021 | 0.0035 |
| RXN-8629 | -0.0021 | 0.0035 |
| RXN0-1132 | -0.0021 | 0.0035 |
| RXN-8001 | 0.0011 | 0.0036 |
| OROPRIBTRANS-RXN | 0.00078 | 0.0036 |
| 6.3.2.10-RXN | 0.00073 | 0.0037 |
| UDP-NACMURALGLDAPAALIG-RXN | 0.00073 | 0.0037 |
| NACGLCTRANS-RXN | 0.00078 | 0.0037 |
| RXN-11029 | 0.00078 | 0.0037 |
| RXN-11346 | 0.00078 | 0.0037 |
| RXN-8976 | 0.00078 | 0.0037 |
| UDPNACETYLGLUCOSAMENOLPYRTRANS-RXN | 0.00073 | 0.0038 |
| RIBOSYLHOMOCYSTEINASE-RXN | 0.0013 | 0.0038 |
| PRPPSYN-RXN | 0.0012 | 0.0040 |
| GLUC1PADENYLTRANS-RXN | 0.0024 | 0.0040 |
| GCVT-RXN | -0.0016 | 0.0041 |
| HEMN-RXN | 0.0010 | 0.0041 |
| GLUTAMATE-SYNTHASE-NADH-RXN | 0.0030 | 0.0043 |
| DIHYDROOROT-RXN | 0.00094 | 0.0043 |
| GLYCOGENSYN-RXN | 0.0015 | 0.0051 |
| RXN-13163 | 0.0015 | 0.0058 |

**Supplementary Table 4:** MetaCyc metabolic pathway comparisons in cases and controls in discovery and validation cohorts, identified by linear modelling.

| Discovery cohort | | |
|---|---|---|
| **Feature** | **Coefficient** | **P-value** |
| PWY_7234_inosine_5__phosphate_biosynthesis_III | -0.0015 | 0.001 |
| PWY_7383_anaerobic_energy_metabolism_invertebrates_cytosol_ | -0.0014 | 0.021 |
| PWY_72824_amino_2_methyl_5_phosphomethyl_pyrimidine_biosynthesis_yeast_ | -0.0012 | 0.036 |
| GLYCOGENSYNTH_PWY_glycogen_biosynthesis_I_from_ADP_D_Glucose_ | 0.0013 | 0.036 |
| HISTSYN_PWY_L_histidine_biosynthesis | -0.00074 | 0.039 |
| PWY_4242_pantothenate_and_coenzymeA_biosynthesis_III | -0.00087 | 0.042 |
| PWY_6700_queuosine_biosynthesis | 0.0012 | 0.044 |
| PWY66_399_gluconeogenesis_III | -0.00094 | 0.045 |

| Validation cohort | | |
|---|---|---|
| **Feature** | **Coefficient** | **P-value** |
| ARO_PWY_chorismate_biosynthesis_I | 0.0025 | 0.0011 |
| PWY_6151S_adenosyl_L_methioninecycle_I | 0.0027 | 0.0012 |
| COMPLETE_ARO_PWY_superpathway_of_aromatic_amino_acid_biosynthesis | 0.0023 | 0.0014 |
| PWY_6163_chorismate_biosynthesis_from_3_dehydroquinate | 0.0027 | 0.0014 |
| PYRIDOXSYN_PWY_pyridoxal_5__phosphatebiosynthesis_I | -0.0026 | 0.011 |
| NONMEVIPP_PWY_methylerythritol_phosphate_pathway_I | 0.0017 | 0.013 |
| NONOXIPENT_PWY_pentose_phosphate_pathway_non_oxidative_branch_ | 0.0024 | 0.022 |
| PWY_6386_UDP_N_acetylmuramoyl_pentapeptide_biosynthesis_II_lysine_containing_ | 0.0017 | 0.029 |
| ARGININE_SYN4_PWY_L_ornithine_denovo_biosynthesis | -0.0027 | 0.031 |
| PWY_6317_galactose_degradation_I_Le_loir_pathway_ | 0.0020 | 0.036 |
| SER_GLYSYN_PWY_superpathway_of_L_serine_and_glycine_biosynthesis_I | 0.0017 | 0.037 |
| PWY66_422_D_galactose_degradation_V_Le_loir_pathway_ | 0.0019 | 0.044 |
| PWY_7357_thiamin_formation_from_pyrithiamine_and_oxythiamine_yeast_ | 0.0017 | 0.045 |
| PWY0_1296_purine_ribonucleosides_degradation | 0.0017 | 0.047 |
| PWY_6387UDP_N_acetylmuramoyl_pentapeptide_biosynthesis_I_meso_diaminopimelate_containing_ | 0.0014 | 0.049 |

**Supplementary Table 5:** Correlation between the indicator species (cases and controls) and the commonly-differentiated KEGG Orthogroups.

| | EC 2.6.1.9: Histidinol-phosphate transaminase | | EC 2.7.4.1: Polyphosphate kinase | | EC 4.3.3.6: Pyridoxal 5'-phosphate synthase | | EC 1.15.1.1: Superoxide proteinase | | EC 3.4.21.53: ATP-dependent serine phosphatase | | EC 2.4.2.17: ATP phosphoribosyltransferase | |
|---|---|---|---|---|---|---|---|---|---|---|---|---|
| | $r^2$ | P-value | $r^2$ | P-value | $r^2$ | P-value | $r^2$ | P-value | $r^2$ | P-value | $r^2$ | P-value |
| Bifidobacterium adolescentis | 0.0078 | 0.012 | 0.012 | 0.00036 | 0.00080 | 0.019 | 0.011 | 0.044 | 0.0092 | 2.63E-04 | 0.0056 | 0.0013 |
| Clostridiales bacterium 1 7 47FAA | 0.056 | 4.04E-06 | 0.054 | 1.49E-05 | 0.017 | 6.71E-04 | 0.027 | 6.68E-05 | 0.033 | 1.47E-05 | 0.028 | 5.72E-04 |
| Clostridium bolteae | 0.068 | 2.62E-06 | 0.099 | 2.90E-07 | 0.052 | 3.34E-07 | 0.131 | 2.12E-06 | 0.10 | 3.18E-09 | 0.048 | 3.25E-05 |
| Clostridium hathewayi | 0.054 | 5.91E-05 | 0.072 | 1.36E-06 | 0.042 | 9.83E-07 | 0.047 | 1.25E-06 | 0.063 | 1.08E-06 | 0.015 | 1.19E-04 |
| Coprococcus comes | 0.032 | 8.06E-04 | 0.076 | 2.31E-07 | 0.0090 | 0.020 | 0.061 | 1.75E-05 | 0.045 | 5.65E-06 | 0.019 | 1.20E-04 |
| Lachnospiraceae bacterium 5 1 63FAA | - | NS | - | NS | - | NS | - | NS | - | NS | - | NS |
| Roseburia inulinivorans | 0.022 | 0.010 | 0.034 | 4.01E-04 | - | NS | 0.040 | 1.66E-03 | - | NS | 0.065 | 6.96E-04 |

**Supplementary Table 6:** Differentially abundant bacterial species according to TNFi treatment, identified by linear modelling.

| Feature | Value | Coefficient | P-value |
|---|---|---|---|
| Bilophila unclassified | Untreated (relative to HC) | -0.0118 | 0.0218 |
| Bilophila unclassified | Untreated (relative to Treated) | -0.0146 | 0.0133 |
| Bilophila unclassified | Treated (relative to HC) | - | NS |
| Clostridium symbiosum | Untreated (relative to HC) | 0.004 | 0.0033 |
| Clostridium symbiosum | Untreated (relative to Treated) | 0.0033 | 0.0366 |
| Clostridium symbiosum | Treated (relative to HC) | - | NS |
| Eggerthella unclassified | Untreated (relative to HC) | 0.0055 | 0.0022 |
| Eggerthella unclassified | Untreated (relative to Treated) | 0.006 | 0.0034 |
| Eggerthella unclassified | Treated (relative to HC) | - | NS |
| Eubacterium biforme | Untreated (relative to HC) | -0.0129 | 0.0275 |
| Eubacterium biforme | Untreated (relative to Treated) | -0.0152 | 0.0238 |
| Eubacterium biforme | Treated (relative to HC) | - | NS |
| Faecalibacterium prausnitzii | Untreated (relative to HC) | -0.0739 | 0.0052 |
| Faecalibacterium prausnitzii | Untreated (relative to Treated) | -0.0751 | 0.0129 |
| Faecalibacterium prausnitzii | Treated (relative to HC) | - | NS |
| Klebsiella pneumoniae | Untreated (relative to HC) | -0.0101 | 0.0153 |
| Klebsiella pneumoniae | Untreated (relative to Treated) | - | NS |
| Klebsiella pneumoniae | Treated (relative to HC) | - | NS |
| Prevotella copri | Untreated (relative to HC) | -0.0518 | 0.0231 |
| Prevotella copri | Untreated (relative to Treated) | - | NS |
| Prevotella copri | Treated (relative to HC) | - | NS |
| Ruminococcus bromii | Untreated (relative to HC) | -0.0318 | 0.0193 |
| Ruminococcus bromii | Untreated (relative to Treated) | -0.0324 | 0.0377 |
| Ruminococcus bromii | Treated (relative to HC) | - | NS |

**Supplementary Table 7:** Differentially abundant MetaCyc metabolic pathways for TNFi treatment, identified by linear modelling. *** P < 0.001, ** P < 0.01, * P < 0.05.

| Class | Pathway | HC vs Treated | HC vs Untreated | Treated vs Untreated |
|---|---|---|---|---|
| Amino acid biosynthesis | BRANCHED-CHAIN-AA-SYN-PWY Superpathway of branched amino acid biosynthesis | | * | ** |
| Amino acid biosynthesis | COMPLETE-ARO-PWY Superpathway of aromatic amino acid biosynthesis | | *** | * |
| Amino acid biosynthesis | ILEUSYN-PWY L-isoleucine biosynthesis (from threonine) | * | * | ** |
| Amino acid biosynthesis | PWY-5103 L-isoleucine biosynthesis (from glutamate) | | * | ** |
| Amino acid biosynthesis | PWY-5347 Superpathway of L-methionine biosynthesis | | * | * |
| Amino acid biosynthesis | VALSYN-PWY L-valine biosynthesis | * | * | ** |
| Aromatic compound biosynthesis | PWY-6163 Chorismate biosynthesis from 3-dehydroquinate | | *** | * |
| Carbohydrate metabolism | CALVIN-PWY Calvin Benson Bassham cycle | | * | ** |
| Carbohydrate metabolism | PWY-6317 Galactose degradation (Leloir pathway) | | * | * |
| Carbohydrate metabolism | PWY-6737 Starch degradation | | * | * |
| Carbohydrate metabolism | PEPTIDOGLYCANSYN-PWY Peptidoglycan biosynthesis | | * | ** |
| Cell structure | PWY-6386 UDP N-acetylmuramoyl pentapeptide biosynthesis (lysine containing) | | * | * |
| Cell structure | NONOXIPENT-PWY Pentose phosphate pathway (non-oxidative branch) | | * | ** |
| Metabolites and energy | PWY-6700 Queuosine biosynthesis | | ** | * |
| Nucleic acid processing | PWY-5686 UMP biosynthesis | * | * | ** |
| Nucleotide biosynthesis | PWY-6545 Pyrimidine deoxyribonucleotides denovo biosynthesis | | * | ** |
| Nucleotide biosynthesis | PWY-7219 Adenosine ribonucleotides denovo biosynthesis | | * | ** |
| Nucleotide biosynthesis | PWY-7221 Guanosine ribonucleotides denovo biosynthesis | | * | ** |
| Nucleotide biosynthesis | PWY-7234 Inosine 5-phosphate biosynthesis | | *** | * |
| Secondary metabolite biosyn. | NONMEVIPP-PWY Methylerythritol phosphate pathway | | * | * |

| Pathway | Value | Coefficient | P-value |
|---|---|---|---|
| BRANCHED-CHAIN-AA-SYN-PWY Superpathway of branched amino acid biosynthesis | Untreated (relative to HC) | -0.0014 | 0.0118 |
| BRANCHED-CHAIN-AA-SYN-PWY Superpathway of branched amino acid biosynthesis | Untreated (relative to Treated) | -0.0022 | 0.0014 |
| COMPLETE-ARO-PWY Superpathway of aromatic amino acid biosynthesis | Untreated (relative to HC) | -0.0015 | 0.0002 |
| COMPLETE-ARO-PWY Superpathway of aromatic amino acid biosynthesis | Untreated (relative to Treated) | -0.0011 | 0.0286 |
| ILEUSYN-PWY L-isoleucine biosynthesis (from threonine) | Untreated (relative to HC) | -0.0010 | 0.0470 |
| ILEUSYN-PWY L-isoleucine biosynthesis (from threonine) | Untreated (relative to Treated) | -0.0019 | 0.0010 |
| ILEUSYN-PWY L-isoleucine biosynthesis (from threonine) | HC (relative to Treated) | -0.0010 | 0.0525 |
| PWY-5103 L-isoleucine biosynthesis (from glutamate) | Untreated (relative to HC) | -0.0014 | 0.0161 |
| PWY-5103 L-isoleucine biosynthesis (from glutamate) | Untreated (relative to Treated) | -0.0022 | 0.0026 |
| PWY-5347 Superpathway of L-methionine biosynthesis | Untreated (relative to HC) | -0.0007 | 0.0480 |
| PWY-5347 Superpathway of L-methionine biosynthesis | Untreated (relative to Treated) | -0.0010 | 0.0290 |
| VALSYN-PWY L-valine biosynthesis | Untreated (relative to HC) | -0.0010 | 0.0470 |
| VALSYN-PWY L-valine biosynthesis | Untreated (relative to Treated) | -0.0019 | 0.0010 |
| VALSYN-PWY L-valine biosynthesis | HC (relative to Treated) | -0.0010 | 0.0525 |
| PWY-6163 Chorismate biosynthesis from 3-dehydroquinate | Untreated (relative to HC) | -0.0018 | 0.0002 |
| PWY-6163 Chorismate biosynthesis from 3-dehydroquinate | Untreated (relative to Treated) | -0.0012 | 0.0356 |
| CALVIN-PWY Calvin Benson Bassham cycle | Untreated (relative to HC) | -0.0011 | 0.0300 |
| CALVIN-PWY Calvin Benson Bassham cycle | Untreated (relative to Treated) | -0.0018 | 0.0028 |
| PEPTIDOGLYCANSYN-PWY Peptidoglycan biosynthesis | Untreated (relative to HC) | -0.0011 | 0.0030 |
| PEPTIDOGLYCANSYN-PWY Peptidoglycan biosynthesis | Untreated (relative to Treated) | -0.0014 | 0.0021 |
| PWY-6317 Galactose degradation (Leloir pathway) | Untreated (relative to HC) | -0.0012 | 0.0412 |
| PWY-6317 Galactose degradation (Leloir pathway) | Untreated (relative to Treated) | -0.0014 | 0.0486 |
| PWY-6737 Starch degradation | Untreated (relative to HC) | -0.0020 | 0.0033 |
| PWY-6737 Starch degradation | Untreated (relative to Treated) | -0.0023 | 0.0076 |
| NONOXIPENT-PWY Pentose phosphate pathway (non-oxidative branch) | Untreated (relative to HC) | -0.0017 | 0.0053 |
| NONOXIPENT-PWY Pentose phosphate pathway (non-oxidative branch) | Untreated (relative to Treated) | -0.0022 | 0.0031 |
| PWY-6386 UDP N-acetylmuramoyl pentapeptide biosynthesis (lysine containing) | Untreated (relative to HC) | -0.0012 | 0.0030 |
| PWY-6386 UDP N-acetylmuramoyl pentapeptide biosynthesis (lysine containing) | Untreated (relative to Treated) | -0.0015 | 0.0041 |
| PWY-6700 Queuosine biosynthesis | Untreated (relative to HC) | -0.0018 | 0.0031 |
| PWY-6700 Queuosine biosynthesis | Untreated (relative to Treated) | -0.0017 | 0.0190 |
| PWY-5686 UMP biosynthesis | Untreated (relative to HC) | -0.0014 | 0.0043 |
| PWY-5686 UMP biosynthesis | Untreated (relative to Treated) | -0.0022 | 0.0002 |
| PWY-5686 UMP biosynthesis | HC (relative to Treated) | -0.0008 | 0.1187 |
| PWY-6545 Pyrimidine deoxyribonucleotides denovo biosynthesis | Untreated (relative to HC) | -0.0011 | 0.0135 |

| | | | |
|---|---|---|---|
| PWY-6545 Pyrimidine deoxyribonucleotides denovo biosynthesis | Untreated (relative to Treated) | -0.0016 | 0.0030 |
| PWY-7219 Adenosine ribonucleotides denovo biosynthesis | Untreated (relative to HC) | -0.0019 | 0.0015 |
| PWY-7219 Adenosine ribonucleotides denovo biosynthesis | Untreated (relative to Treated) | -0.0023 | 0.0016 |
| PWY-7221 Guanosine ribonucleotides denovo biosynthesis | Untreated (relative to HC) | -0.0013 | 0.0009 |
| PWY-7221 Guanosine ribonucleotides denovo biosynthesis | Untreated (relative to Treated) | -0.0018 | 0.0002 |
| PWY-7234 Inosine 5-phosphate biosynthesis | Untreated (relative to HC) | 0.0017 | 0.0003 |
| PWY-7234 Inosine 5-phosphate biosynthesis | Untreated (relative to Treated) | 0.0013 | 0.0242 |
| NONMEVIPP-PWY Methylerythritol phosphate pathway | Untreated (relative to HC) | -0.0011 | 0.0033 |
| NONMEVIPP-PWY Methylerythritol phosphate pathway | Untreated (relative to Treated) | -0.0013 | 0.0048 |

**Supplementary Table 8:** Correlation between differentially abundant bacterial species and MetaCyc metabolic pathways for TNFi treatment.

| | BRANCHED-CHAIN-AA-SYN-PWY: Superpathway of branched amino acid biosynthesis | | COMPLETE-ARO-PWY: Superpathway of aromatic amino acid biosynthesis | | ILEUSYN-PWY: L-isoleucine biosynthesis (from threonine) | | PWY-5103: L-isoleucine biosynthesis (from glutamate) | | PWY-5347: Superpathway of L-methionine biosynthesis (transsulfuration) | | VALSYN-PWY: L-valine biosynthesis | | PWY-6163: Chorismate biosynthesis from 3-dehydroquinate | |
|---|---|---|---|---|---|---|---|---|---|---|---|---|---|---|
| | $r^2$ | P-value | $r^2$ | P-value | $r^2$ | P-value | $r^2$ | P-value | $r^2$ | P-value | $r^2$ | P-value | $r^2$ | P-value |
| Prevotella copri | - | NS | - | NS | - | NS | - | NS | - | NS | - | NS | - | NS |
| Faecalibacterium prausnitzii | 0.387 | 6.18E-28 | 0.027 | 1.33E-03 | 0.292 | 2.19E-21 | 0.3870 | 3.46E-27 | - | NS | 0.2922 | 2.19E-21 | 0.028 | 1.52E-03 |
| Bilophila unclassified | - | NS | -0.030 | 6.22E-03 | - | NS | - | NS | - | NS | - | NS | -0.031 | 5.00E-03 |
| Klebsiella pneumoniae | - | NS | -0.025 | 3.29E-02 | -0.034 | 5.05E-03 | - | NS | - | NS | -0.034 | 5.05E-03 | - | NS |
| Ruminococcus bromii | - | NS | 0.031 | 5.79E-03 | 0.049 | 1.13E-03 | - | NS | 0.218 | 6.55E-05 | 0.049 | 1.13E-03 | 0.035 | 2.55E-03 |
| Eubacterium biforme | - | NS | 0.002 | 1.17E-02 | - | NS | - | NS | 0.021 | 4.51E-02 | - | NS | 0.003 | 8.83E-03 |
| Clostridium symbiosum | 0.014 | 2.43E-05 | 0.015 | 7.62E-05 | 0.004 | 3.01E-02 | 0.011 | 2.59E-05 | - | NS | 0.004 | 3.01E-02 | 0.018 | 8.00E-05 |
| Eggerthella unclassified | - | NS | 0.018 | 2.91E-03 | - | NS | - | NS | - | NS | - | NS | 0.022 | 1.24E-03 |

| | PEPTIDOGLYCANSYN-PWY: Peptidoglycan biosynthesis (meso-diaminopimelate containing) | | PWY-6386: UDP-N-acetylmuramoyl pentapeptide biosynthesis (lysine containing) | | NONOXIPENT-PWY: Pentose phosphate pathway (non-oxidative branch) | | PWY-6700: Queuosine biosynthesis | | PWY-5686: UMP biosynthesis | | PWY-6545: Pyrimidine deoxyribonucleotides denovo biosynthesis | | PWY-7219: Adenosine ribonucleotides de novo biosynthesis | |
|---|---|---|---|---|---|---|---|---|---|---|---|---|---|---|
| | $r^2$ | P-value | $r^2$ | P-value | $r^2$ | P-value | $r^2$ | P-value | $r^2$ | P-value | $r^2$ | P-value | $r^2$ | P-value |
| Prevotella copri | 0.104 | 3.97E-03 | 0.055 | 1.32E-02 | - | NS | 0.3243 | 5.40E-03 | 0.115 | 9.18E-03 | - | NS | 0.126 | 2.53E-03 |
| Faecalibacterium prausnitzii | 0.067 | 4.27E-07 | 0.104 | 1.13E-09 | 0.089 | 3.43E-09 | 0.030 | 7.50E-13 | 0.107 | 6.00E-12 | - | NS | 0.070 | 4.65E-09 |
| Bilophila unclassified | - | NS | - | NS | -0.017 | 1.57E-02 | - | NS | - | NS | - | NS | - | NS |
| Klebsiella pneumoniae | -0.045 | 3.70E-03 | -0.044 | 2.00E-03 | - | NS | -0.003 | 4.89E-03 | -0.043 | 9.18E-03 | - | NS | -0.032 | 2.61E-02 |
| Ruminococcus bromii | - | NS | 0.033 | 4.23E-02 | 0.053 | 4.35E-03 | 0.008 | 4.27E-02 | 0.025 | 4.88E-02 | - | NS | - | NS |
| Eubacterium biforme | - | NS | - | NS | - | NS | 0.000 | 5.74E-03 | 0.010 | 9.18E-03 | - | NS | 0.016 | 7.61E-04 |
| Clostridium symbiosum | 0.011 | 1.18E-02 | 0.016 | 2.10E-03 | 0.009 | 8.51E-03 | 0.010 | 1.34E-03 | 0.015 | 5.36E-03 | - | NS | 0.036 | 5.82E-05 |
| Eggerthella unclassified | - | NS | 0.012 | 1.98E-02 | - | NS | 0.013 | 7.32E-03 | 0.011 | 9.76E-03 | - | NS | 0.017 | 8.20E-04 |

| | CALVIN-PWY: Calvin Benson Bassham cycle | | PWY-6317: Galactose degradation (Leloir pathway) | | PWY-6737: Starch degradation | | PWY-7221: Guanosine ribonucleotides de novo biosynthesis | | PWY-7234: Inosine-5-phosphate biosynthesis | | NONMEVIPP-PWY: Methylerythritol phosphate pathway | |
|---|---|---|---|---|---|---|---|---|---|---|---|---|
| | **$r^2$** | **P-value** | **$r^2$** | **P-value** | **$r^2$** | **P-value** | **$r^2$** | **P-value** | **$r^2$** | **P-value** | **$r^2$** | **P-value** |
| Prevotella copri | - | NS | - | NS | - | NS | 0.2923 | 1.88E-02 | 0.039 | 7.68E-04 | - | NS |
| Faecalibacterium prausnitzii | 0.044 | 3.60E-08 | 0.076 | 1.78E-11 | 0.312 | 2.38E-24 | 0.042 | 1.06E-09 | 0.012 | 2.89E-02 | 0.040 | 6.18E-28 |
| Bilophila unclassified | - | NS | -0.029 | 3.60E-03 | - | NS | - | NS | - | NS | - | NS |
| Klebsiella pneumoniae | -0.014 | 2.94E-02 | - | NS | -0.043 | 1.41E-02 | - | NS | - | NS | - | NS |
| Ruminococcus bromii | 0.032 | 1.19E-02 | 0.029 | 1.68E-04 | 0.034 | 1.35E-04 | 0.031 | 3.15E-03 | - | NS | - | NS |
| Eubacterium biforme | - | NS | - | NS | - | NS | - | NS | 0.016 | 1.07E-03 | - | NS |
| Clostridium symbiosum | 0.007 | 1.50E-02 | 0.013 | 2.34E-04 | 0.034 | 5.86E-06 | - | NS | 0.057 | 3.13E-03 | 0.023 | 2.43E-05 |
| Eggerthella unclassified | - | NS | - | NS | - | NS | - | NS | 0.018 | 2.75E-03 | - | NS |

**Supplementary Table 9:** Differentially abundant species for *RUNX3* variants in the Han-Chinese cohort, identified by linear modelling. Differential abundance is calculated relative to the AA genotype.

| Variable | Feature | Value | Coefficient | P-value |
|---|---|---|---|---|
| RUNX3 | Bacteroides caccae | AG | -0.0176 | 0.0450 |
| RUNX3 | Bacteroides coprocola | AG | -0.0359 | 0.0207 |
| RUNX3 | Bacteroides massiliensis | AG | -0.0175 | 0.0482 |
| RUNX3 | Bacteroides plebeius | AG | -0.0320 | 0.0174 |
| RUNX3 | Bacteroides stercoris | AG | -0.0376 | 0.0150 |
| RUNX3 | Bacteroides uniformis | AG | -0.0395 | 0.0029 |
| RUNX3 | Bacteroides uniformis | GG | -0.0399 | 0.0158 |
| RUNX3 | Barnesiella intestinihominis | AG | -0.0104 | 0.0452 |
| RUNX3 | Burkholderiales bacterium 1 1 47 | AG | -0.0083 | 0.0231 |
| RUNX3 | Citrobacter freundii | GG | -0.0080 | 0.0467 |
| RUNX3 | Citrobacter unclassified | GG | -0.0079 | 0.0393 |
| RUNX3 | Dorea formicigenerans | AG | -0.0130 | 0.0390 |
| RUNX3 | Eubacterium ramulus | AG | -0.0094 | 0.0416 |
| RUNX3 | Eubacterium ventriosum | AG | -0.0315 | 0.0000 |
| RUNX3 | Eubacterium ventriosum | GG | -0.0268 | 0.0054 |
| RUNX3 | Lachnospiraceae bacterium 1 1 57FAA | AG | -0.0255 | 0.0035 |
| RUNX3 | Lachnospiraceae bacterium 1 1 57FAA | GG | -0.0237 | 0.0298 |
| RUNX3 | Pantoea unclassified | AG | 0.0077 | 0.0383 |
| RUNX3 | Parabacteroides merdae | AG | -0.0236 | 0.0072 |
| RUNX3 | Rothia mucilaginosa | AG | -0.0071 | 0.0021 |
| RUNX3 | Rothia mucilaginosa | GG | -0.0065 | 0.0240 |
| RUNX3 | Streptococcus australis | AG | -0.0032 | 0.0110 |
| RUNX3 | Streptococcus australis | GG | -0.0033 | 0.0362 |
| RUNX3 | Streptococcus parasanguinis | AG | -0.0075 | 0.0402 |
| RUNX3 | Subdoligranulum unclassified | AG | -0.0360 | 0.0206 |

**Supplementary Table 10:** Differentially abundant MetaCyc metabolic pathways for *RUNX3* variants in the Han-Chinese cohort, identified by linear modelling. Differential abundance is calculated relative to the AA genotype.

| Variable | Feature | Value | Coefficient | P-value |
|---|---|---|---|---|
| RUNX3 | PANTOSYN-PWY: pantothenate and coenzyme A biosynthesis I | AG | -0.0012 | 0.0006 |
| RUNX3 | COA-PWY: coenzyme A biosynthesis I | AG | -0.0014 | 0.0036 |
| RUNX3 | PWY-5659:GDP mannose biosynthesis | AG | -0.0015 | 0.0056 |
| RUNX3 | PWY-6737: starch degradation V | AG | -0.0020 | 0.0070 |
| RUNX3 | GLYCOLYSIS: glycolysis I from glucose 6 phosphate | AG | 0.0015 | 0.0070 |
| RUNX3 | PWY-7187: pyrimidine deoxyribonucleotides denovo biosynthesis II | GG | -0.0009 | 0.0071 |
| RUNX3 | THRESYN-PWY: superpathway of L threonine biosynthesis | AG | -0.0009 | 0.0082 |
| RUNX3 | PWY-5484: glycolysis II from fructose 6 phosphate | AG | 0.0014 | 0.0092 |
| RUNX3 | PWY-5100: pyruvate fermentation to acetate and lactate II | AG | -0.0014 | 0.0110 |
| RUNX3 | PWY-7229: superpathway of fadenosine nucleotides denovo biosynthesis I | GG | -0.0012 | 0.0110 |
| RUNX3 | POLYAMSYN-PWY: superpathway of polyamine biosynthesis I | GG | -0.0014 | 0.0136 |
| RUNX3 | PWY-3001: superpathway of L isoleucine biosynthesis I | AG | -0.0010 | 0.0138 |
| RUNX3 | PWY-5104: L isoleucine biosynthesis IV | AG | -0.0019 | 0.0161 |
| RUNX3 | PWY-3001: superpathway of L isoleucine biosynthesis I | GG | -0.0012 | 0.0170 |
| RUNX3 | PWY-61215: aminoimidazole ribonucleotide biosynthesis I | AG | -0.0010 | 0.0174 |
| RUNX3 | PWY-6126: superpathway of adenosine nucleotides denovo biosynthesis II | GG | -0.0011 | 0.0186 |
| RUNX3 | ANAEROFRUCAT-PWY: homolactic fermentation | GG | -0.0012 | 0.0230 |
| RUNX3 | PWY-6317: galactose degradation I Le loir pathway | AG | -0.0014 | 0.0245 |
| RUNX3 | PWY0-166: superpathway of pyrimidine deoxyribonucleotides denovo biosynthesis E coli | GG | -0.0011 | 0.0284 |
| RUNX3 | THRESYN-PWY: superpathway of L threoninebiosynthesis | GG | -0.0010 | 0.0288 |
| RUNX3 | PWY66-422: D galactose degradation V Le loir pathway | AG | -0.0014 | 0.0296 |
| RUNX3 | GALACTUROCAT-PWY: D galacturonate degradation I | AG | -0.0010 | 0.0323 |
| RUNX3 | PWY0-162: superpathway of pyrimidine ribonucleotides denovo biosynthesis | GG | -0.0021 | 0.0339 |
| RUNX3 | PWY-1042: glycolysis IV plant cytosol | GG | -0.0016 | 0.0371 |
| RUNX3 | PWY-5177: glutaryl CoA degradation | AG | -0.0012 | 0.0398 |
| RUNX3 | PWY-1042: glycolysis IV plantcytosol | AG | -0.0012 | 0.0404 |
| RUNX3 | PWY-5100: pyruvate fermentation to acetate and lactate II | GG | -0.0014 | 0.0423 |
| RUNX3 | ILEUSYN-PWY: L isoleucine biosynthesis I from threonine | AG | -0.0011 | 0.0431 |

**Supplementary Table 11:** Correlation between the indicator species and the MetaCyc metabolic pathways.

| | **POLYAMSYN-PWY: Superpathway of polyamine biosynthesis** | | **THRESYN-PWY: Superpathway of L-threonine biosynthesis** | | **PWY-3001: Superpathway of L-isoleucine biosynthesis** | | **PWY-7229: Superpathway of adenosine nucleotides denovo biosynthesis** | | **PWY-6126: Superpathway of adenosine nucleotides denovo biosynthesis** | | **PWY-5100: Pyruvate fermentation to acetate and lactate** | | **PWY-7187: Pyrimidine deoxyribonucleotides denovo biosynthesis** | |
|---|---|---|---|---|---|---|---|---|---|---|---|---|---|---|
| | **$r^2$** | **P-value** | **$r^2$** | **P-value** | **$r^2$** | **P-value** | **$r^2$** | **P-value** | **$r^2$** | **P-value** | **$r^2$** | **P-value** | **$r^2$** | **P-value** |
| Lachnospiraceae bacterium 1_1_57FAA | - | NS | - | NS | - | NS | - | NS | 0.0049 | 0.0429 | 0.0082 | 0.0450 | - | NS |
| Eubacterium ventriosum | - | NS | 0.0148 | 0.0131 | 0.0210 | 0.0034 | - | NS | - | NS | 0.0715 | 0.0000 | - | NS |
| Citrobacter unclassified | - | NS | - | NS | - | NS | - | NS | - | NS | - | NS | - | NS |
| Citrobacter freundii | - | NS | - | NS | - | NS | - | NS | - | NS | - | NS | - | NS |

**Supplementary Table 12:** Differentially abundant species for *RUNX3* variants, as detected by 16S amplicon sequencing on a predominately Caucasian cohort, identified by linear modelling. Subjects were sampled from six sites: rectum, cecum, ileum, right colon (RC), left colon (LC) and stool. Differential abundance is calculated relative to the AA genotype.

| Variable | Feature | Value | Coefficient | P-value | Site |
|---|---|---|---|---|---|
| RUNX3 | k__Bacteria_p__Actinobacteria_c__Actinobacteria_o__Actinomycetales_f__Actinomycetaceae_g__Varibaculum | GG | -0.0164 | 0.0266 | Rectum |
| RUNX3 | k__Bacteria_p__Actinobacteria_c__Actinobacteria_o__Actinomycetales_f__Actinomycetaceae_g__Varibaculum | AG | -0.0152 | 0.0278 | Rectum |
| RUNX3 | k__Bacteria_p__Actinobacteria_c__Actinobacteria_o__Actinomycetales_f__Actinomycetaceae_g__Varibaculum_s__ | GG | -0.0164 | 0.0266 | Rectum |
| RUNX3 | k__Bacteria_p__Actinobacteria_c__Actinobacteria_o__Actinomycetales_f__Actinomycetaceae_g__Varibaculum_s__ | AG | -0.0152 | 0.0278 | Rectum |
| RUNX3 | k__Bacteria_p__Actinobacteria_c__Actinobacteria_o__Bifidobacteriales_f__Bifidobacteriaceae_g__Bifidobacterium_s__longum | GG | 0.0057 | 0.0207 | Cecum |
| RUNX3 | k__Bacteria_p__Bacteroidetes_c__Bacteroidia_o__Bacteroidales_f__Bacteroidaceae_g__Bacteroides_s__ovatus | GG | 0.0400 | 0.0131 | Stool |
| RUNX3 | k__Bacteria_p__Bacteroidetes_c__Bacteroidia_o__Bacteroidales_f__Bacteroidaceae_g__Bacteroides_s__ovatus | AG | 0.0354 | 0.0198 | Stool |
| RUNX3 | k__Bacteria_p__Bacteroidetes_c__Bacteroidia_o__Bacteroidales_f__Bacteroidaceae_g__Bacteroides_s__ovatus | AG | 0.0272 | 0.0399 | Cecum |
| RUNX3 | k__Bacteria_p__Bacteroidetes_c__Bacteroidia_o__Bacteroidales_f__Prevotellaceae | AG | -0.0327 | 0.0260 | Cecum |
| RUNX3 | k__Bacteria_p__Bacteroidetes_c__Bacteroidia_o__Bacteroidales_f__Prevotellaceae | AG | -0.0099 | 0.0276 | Ileum |
| RUNX3 | k__Bacteria_p__Bacteroidetes_c__Bacteroidia_o__Bacteroidales_f__Prevotellaceae_g__Prevotella | AG | -0.0327 | 0.0260 | Cecum |
| RUNX3 | k__Bacteria_p__Bacteroidetes_c__Bacteroidia_o__Bacteroidales_f__Prevotellaceae_g__Prevotella | AG | -0.0099 | 0.0276 | Ileum |
| RUNX3 | k__Bacteria_p__Bacteroidetes_c__Bacteroidia_o__Bacteroidales_f__Prevotellaceae_g__Prevotella_s__ | AG | -0.0243 | 0.0062 | Rectum |
| RUNX3 | k__Bacteria_p__Bacteroidetes_c__Bacteroidia_o__Bacteroidales_f__Prevotellaceae_g__Prevotella_s__ | GG | -0.0214 | 0.0233 | Rectum |
| RUNX3 | k__Bacteria_p__Bacteroidetes_c__Bacteroidia_o__Bacteroidales_f__Prevotellaceae_g__Prevotella_s__stercorea | AG | -0.0048 | 0.0305 | Cecum |
| RUNX3 | k__Bacteria_p__Bacteroidetes_c__Bacteroidia_o__Bacteroidales_f__Prevotellaceae_g__Prevotella_s__stercorea | GG | -0.0048 | 0.0388 | Cecum |
| RUNX3 | k__Bacteria_p__Bacteroidetes_c__Bacteroidia_o__Bacteroidales_f__Prevotellaceae_g__Prevotella_s__stercorea | AG | -0.0612 | 0.0432 | Ileum |
| RUNX3 | k__Bacteria_p__Bacteroidetes_c__Bacteroidia_o__Bacteroidales_f__S24_7 | AG | 0.0289 | 0.0295 | LC |
| RUNX3 | k__Bacteria_p__Bacteroidetes_c__Bacteroidia_o__Bacteroidales_f__S24_7_g__ | AG | 0.0289 | 0.0295 | LC |
| RUNX3 | k__Bacteria_p__Bacteroidetes_c__Bacteroidia_o__Bacteroidales_f__S24_7_g___s__ | AG | 0.0289 | 0.0295 | LC |
| RUNX3 | k__Bacteria_p__Firmicutes_c__Bacilli_o__Bacillales | AG | -0.0304 | 0.0252 | LC |
| RUNX3 | k__Bacteria_p__Firmicutes_c__Bacilli_o__Bacillales | AG | -0.0697 | 0.0304 | Rectum |
| RUNX3 | k__Bacteria_p__Firmicutes_c__Bacilli_o__Bacillales_f__Paenibacillaceae | AG | -0.0281 | 0.0354 | LC |
| RUNX3 | k__Bacteria_p__Firmicutes_c__Bacilli_o__Bacillales_f__Paenibacillaceae | AG | -0.0623 | 0.0414 | Rectum |
| RUNX3 | k__Bacteria_p__Firmicutes_c__Bacilli_o__Bacillales_f__Paenibacillaceae_g__Paenibacillus | AG | -0.0281 | 0.0354 | LC |
| RUNX3 | k__Bacteria_p__Firmicutes_c__Bacilli_o__Bacillales_f__Paenibacillaceae_g__Paenibacillus | AG | -0.0623 | 0.0414 | Rectum |
| RUNX3 | k__Bacteria_p__Firmicutes_c__Bacilli_o__Bacillales_f__Paenibacillaceae_g__Paenibacillus_s__ | AG | -0.0281 | 0.0354 | LC |
| RUNX3 | k__Bacteria_p__Firmicutes_c__Bacilli_o__Bacillales_f__Paenibacillaceae_g__Paenibacillus_s__ | AG | -0.0623 | 0.0414 | Rectum |
| RUNX3 | k__Bacteria_p__Firmicutes_c__Bacilli_o__Bacillales_f__Staphylococcaceae | AG | -0.0201 | 0.0093 | Rectum |

| | | | | | |
|---|---|---|---|---|---|
| RUNX3 | k__Bacteria_p__Firmicutes_c__Bacilli_o__Bacillales_f__Staphylococcaceae | GG | -0.0199 | 0.0162 | Rectum |
| RUNX3 | k__Bacteria_p__Firmicutes_c__Bacilli_o__Bacillales_f__Staphylococcaceae_g__Staphylococcus | AG | -0.0201 | 0.0093 | Rectum |
| RUNX3 | k__Bacteria_p__Firmicutes_c__Bacilli_o__Bacillales_f__Staphylococcaceae_g__Staphylococcus | GG | -0.0199 | 0.0162 | Rectum |
| RUNX3 | k__Bacteria_p__Firmicutes_c__Bacilli_o__Bacillales_f__Staphylococcaceae_g__Staphylococcus_s__ | AG | -0.0201 | 0.0093 | Rectum |
| RUNX3 | k__Bacteria_p__Firmicutes_c__Bacilli_o__Bacillales_f__Staphylococcaceae_g__Staphylococcus_s__ | GG | -0.0199 | 0.0162 | Rectum |
| RUNX3 | k__Bacteria_p__Firmicutes_c__Bacilli_o__Lactobacillales_f__Lactobacillaceae | AG | -0.0243 | 0.0345 | Rectum |
| RUNX3 | k__Bacteria_p__Firmicutes_c__Bacilli_o__Lactobacillales_f__Lactobacillaceae_g__Lactobacillus | AG | -0.0243 | 0.0345 | Rectum |
| RUNX3 | k__Bacteria_p__Firmicutes_c__Bacilli_o__Lactobacillales_f__Lactobacillaceae_g__Lactobacillus_s__ | AG | -0.0243 | 0.0345 | Rectum |
| RUNX3 | k__Bacteria_p__Firmicutes_c__Clostridia_o__Clostridiales_f__Clostridiaceae | AG | -0.0336 | 0.0013 | RC |
| RUNX3 | k__Bacteria_p__Firmicutes_c__Clostridia_o__Clostridiales_f__Clostridiaceae | AG | -0.0257 | 0.0068 | Ileum |
| RUNX3 | k__Bacteria_p__Firmicutes_c__Clostridia_o__Clostridiales_f__Clostridiaceae_g__ | AG | -0.0161 | 0.0349 | Rectum |
| RUNX3 | k__Bacteria_p__Firmicutes_c__Clostridia_o__Clostridiales_f__Clostridiaceae_g___s__ | AG | -0.0161 | 0.0349 | Rectum |
| RUNX3 | k__Bacteria_p__Firmicutes_c__Clostridia_o__Clostridiales_f__Clostridiaceae_g__SMB53 | AG | -0.0073 | 0.0478 | Ileum |
| RUNX3 | k__Bacteria_p__Firmicutes_c__Clostridia_o__Clostridiales_f__Clostridiaceae_g__SMB53_s__ | AG | -0.0073 | 0.0478 | Ileum |
| RUNX3 | k__Bacteria_p__Firmicutes_c__Clostridia_o__Clostridiales_f__Lachnospiraceae_g___Ruminococcus__s__gnavus | GG | -0.0172 | 0.0216 | LC |
| RUNX3 | k__Bacteria_p__Firmicutes_c__Clostridia_o__Clostridiales_f__Lachnospiraceae_g__Anaerostipes | AG | -0.0102 | 0.0227 | LC |
| RUNX3 | k__Bacteria_p__Firmicutes_c__Clostridia_o__Clostridiales_f__Lachnospiraceae_g__Anaerostipes_s__ | AG | -0.0102 | 0.0227 | LC |
| RUNX3 | k__Bacteria_p__Firmicutes_c__Clostridia_o__Clostridiales_f__Lachnospiraceae_g__Dorea | GG | 0.0331 | 0.0460 | Ileum |
| RUNX3 | k__Bacteria_p__Firmicutes_c__Clostridia_o__Clostridiales_f__Lachnospiraceae_g__Dorea_s__ | GG | 0.0347 | 0.0176 | Rectum |
| RUNX3 | k__Bacteria_p__Firmicutes_c__Clostridia_o__Clostridiales_f__Lachnospiraceae_g__Roseburia | GG | -0.0379 | 0.0357 | Stool |
| RUNX3 | k__Bacteria_p__Firmicutes_c__Clostridia_o__Clostridiales_f__Lachnospiraceae_g__Roseburia_s__ | GG | -0.0379 | 0.0357 | Stool |
| RUNX3 | k__Bacteria_p__Firmicutes_c__Clostridia_o__Clostridiales_f__Peptococcaceae | GG | 0.0077 | 0.0334 | Cecum |
| RUNX3 | k__Bacteria_p__Firmicutes_c__Clostridia_o__Clostridiales_f__Peptococcaceae | AG | 0.0147 | 0.0363 | LC |
| RUNX3 | k__Bacteria_p__Firmicutes_c__Clostridia_o__Clostridiales_f__Ruminococcaceae_g__Oscillospira | GG | -0.0347 | 0.0017 | LC |
| RUNX3 | k__Bacteria_p__Firmicutes_c__Clostridia_o__Clostridiales_f__Ruminococcaceae_g__Oscillospira | AG | -0.0333 | 0.0017 | LC |
| RUNX3 | k__Bacteria_p__Firmicutes_c__Clostridia_o__Clostridiales_f__Ruminococcaceae_g__Oscillospira | GG | -0.0291 | 0.0097 | Cecum |
| RUNX3 | k__Bacteria_p__Firmicutes_c__Clostridia_o__Clostridiales_f__Ruminococcaceae_g__Oscillospira_s__ | GG | -0.0347 | 0.0017 | LC |
| RUNX3 | k__Bacteria_p__Firmicutes_c__Clostridia_o__Clostridiales_f__Ruminococcaceae_g__Oscillospira_s__ | AG | -0.0333 | 0.0017 | LC |
| RUNX3 | k__Bacteria_p__Firmicutes_c__Clostridia_o__Clostridiales_f__Ruminococcaceae_g__Oscillospira_s__ | GG | -0.0291 | 0.0097 | Cecum |
| RUNX3 | k__Bacteria_p__Fusobacteria_c__Fusobacteriia | GG | -0.0194 | 0.0037 | Rectum |
| RUNX3 | k__Bacteria_p__Fusobacteria_c__Fusobacteriia | AG | -0.0130 | 0.0342 | Rectum |
| RUNX3 | k__Bacteria_p__Fusobacteria_c__Fusobacteriia_o__Fusobacteriales | GG | -0.0194 | 0.0037 | Rectum |
| RUNX3 | k__Bacteria_p__Fusobacteria_c__Fusobacteriia_o__Fusobacteriales | AG | -0.0130 | 0.0342 | Rectum |
| RUNX3 | k__Bacteria_p__Fusobacteria_c__Fusobacteriia_o__Fusobacteriales_f__Fusobacteriaceae | GG | -0.0194 | 0.0037 | Rectum |

| | | | | | |
|---|---|---|---|---|---|
| RUNX3 | k__Bacteria_p__Fusobacteria_c__Fusobacteriia_o__Fusobacteriales_f__Fusobacteriaceae | AG | -0.0132 | 0.0323 | Rectum |
| RUNX3 | k__Bacteria_p__Fusobacteria_c__Fusobacteriia_o__Fusobacteriales_f__Fusobacteriaceae_g__Fusobacterium | GG | -0.0194 | 0.0037 | Rectum |
| RUNX3 | k__Bacteria_p__Fusobacteria_c__Fusobacteriia_o__Fusobacteriales_f__Fusobacteriaceae_g__Fusobacterium | AG | -0.0132 | 0.0323 | Rectum |
| RUNX3 | k__Bacteria_p__Fusobacteria_c__Fusobacteriia_o__Fusobacteriales_f__Fusobacteriaceae_g__Fusobacterium_s__ | GG | -0.0194 | 0.0037 | Rectum |
| RUNX3 | k__Bacteria_p__Fusobacteria_c__Fusobacteriia_o__Fusobacteriales_f__Fusobacteriaceae_g__Fusobacterium_s__ | AG | -0.0132 | 0.0323 | Rectum |
| RUNX3 | k__Bacteria_p__Proteobacteria_c__Betaproteobacteria | GG | -0.0492 | 0.0270 | LC |
| RUNX3 | k__Bacteria_p__Proteobacteria_c__Betaproteobacteria | AG | -0.0446 | 0.0369 | LC |
| RUNX3 | k__Bacteria_p__Proteobacteria_c__Betaproteobacteria_o__Burkholderiales | GG | -0.0493 | 0.0244 | LC |
| RUNX3 | k__Bacteria_p__Proteobacteria_c__Betaproteobacteria_o__Burkholderiales | AG | -0.0475 | 0.0244 | LC |
| RUNX3 | k__Bacteria_p__Proteobacteria_c__Betaproteobacteria_o__Burkholderiales_f__ | AG | 0.0026 | 0.0462 | Stool |
| RUNX3 | k__Bacteria_p__Proteobacteria_c__Betaproteobacteria_o__Burkholderiales_f___g__ | AG | 0.0026 | 0.0462 | Stool |
| RUNX3 | k__Bacteria_p__Proteobacteria_c__Betaproteobacteria_o__Burkholderiales_f___g___s__ | AG | 0.0026 | 0.0462 | Stool |
| RUNX3 | k__Bacteria_p__Proteobacteria_c__Betaproteobacteria_o__Burkholderiales_f__Alcaligenaceae | AG | -0.0470 | 0.0312 | LC |
| RUNX3 | k__Bacteria_p__Proteobacteria_c__Betaproteobacteria_o__Burkholderiales_f__Alcaligenaceae | GG | -0.0446 | 0.0483 | LC |
| RUNX3 | k__Bacteria_p__Proteobacteria_c__Betaproteobacteria_o__Burkholderiales_f__Alcaligenaceae_g__Sutterella | AG | -0.0470 | 0.0312 | LC |
| RUNX3 | k__Bacteria_p__Proteobacteria_c__Betaproteobacteria_o__Burkholderiales_f__Alcaligenaceae_g__Sutterella | GG | -0.0446 | 0.0483 | LC |
| RUNX3 | k__Bacteria_p__Proteobacteria_c__Betaproteobacteria_o__Burkholderiales_f__Alcaligenaceae_g__Sutterella_s__ | AG | -0.0470 | 0.0312 | LC |
| RUNX3 | k__Bacteria_p__Proteobacteria_c__Betaproteobacteria_o__Burkholderiales_f__Alcaligenaceae_g__Sutterella_s__ | GG | -0.0446 | 0.0483 | LC |

**Supplementary Table 13:** Differentially abundant KEGG metabolic pathways for *RUNX3* variants, as detected by 16S amplicon sequencing on a predominately Caucasian cohort, identified by linear modelling. Subjects were sampled from six sites: rectum, cecum, ileum, right colon (RC), left colon (LC) and stool. Differential abundance is calculated relative to the AA genotype.

| Variable | Feature | Value | Coefficient | P-value | Site |
|---|---|---|---|---|---|
| RUNX3 | ko00010 | AG | -0.0009 | 0.0436 | RC |
| RUNX3 | ko00240 | AG | -0.0015 | 0.0239 | RC |
| RUNX3 | ko00270 | AG | -0.0011 | 0.0389 | RC |
| RUNX3 | ko00310 | AG | -0.0018 | 0.0069 | Ileum |
| RUNX3 | ko00310 | GG | -0.0014 | 0.0388 | Ileum |
| RUNX3 | ko00311 | GG | -0.0046 | 0.0185 | Ileum |
| RUNX3 | ko00330 | GG | -0.0010 | 0.0369 | LC |
| RUNX3 | ko00350 | AG | -0.0011 | 0.0008 | Ileum |
| RUNX3 | ko00350 | AG | -0.0012 | 0.0115 | RC |
| RUNX3 | ko00350 | AG | -0.0012 | 0.0155 | Rectum |
| RUNX3 | ko00350 | GG | -0.0008 | 0.0264 | Ileum |
| RUNX3 | ko00360 | GG | -0.0020 | 0.0036 | Rectum |
| RUNX3 | ko00440 | GG | -0.0026 | 0.0476 | Stool |
| RUNX3 | ko00471 | AG | -0.0031 | 0.0380 | Cecum |
| RUNX3 | ko00520 | GG | 0.0014 | 0.0275 | Stool |
| RUNX3 | ko00550 | AG | -0.0022 | 0.0269 | RC |
| RUNX3 | ko00564 | GG | 0.0023 | 0.0200 | Ileum |
| RUNX3 | ko00621 | GG | 0.0158 | 0.0197 | Rectum |
| RUNX3 | ko00625 | GG | 0.0208 | 0.0408 | Rectum |
| RUNX3 | ko00626 | AG | -0.0018 | 0.0269 | Rectum |
| RUNX3 | ko00630 | GG | -0.0016 | 0.0082 | Rectum |
| RUNX3 | ko00780 | GG | -0.0026 | 0.0396 | Cecum |
| RUNX3 | ko00780 | GG | -0.0030 | 0.0421 | Ileum |
| RUNX3 | ko00790 | GG | 0.0037 | 0.0476 | Stool |
| RUNX3 | ko00830 | GG | -0.0035 | 0.0073 | Rectum |
| RUNX3 | ko00830 | GG | -0.0033 | 0.0190 | Ileum |
| RUNX3 | ko00830 | GG | -0.0026 | 0.0408 | RC |
| RUNX3 | ko00910 | GG | -0.0013 | 0.0108 | Rectum |
| RUNX3 | ko00941 | AG | -0.0027 | 0.0013 | Rectum |
| RUNX3 | ko00941 | GG | -0.0024 | 0.0119 | Stool |
| RUNX3 | ko00970 | AG | -0.0031 | 0.0314 | RC |
| RUNX3 | ko01040 | AG | -0.0019 | 0.0404 | Ileum |
| RUNX3 | ko02060 | AG | 0.0081 | 0.0350 | Ileum |
| RUNX3 | ko03010 | AG | -0.0027 | 0.0356 | RC |
| RUNX3 | ko03010 | AG | -0.0026 | 0.0410 | Cecum |
| RUNX3 | ko03013 | GG | 0.0011 | 0.0288 | Ileum |
| RUNX3 | ko03070 | AG | -0.0018 | 0.0015 | Ileum |
| RUNX3 | ko03070 | GG | -0.0016 | 0.0096 | Ileum |
| RUNX3 | ko03070 | AG | -0.0014 | 0.0406 | RC |
| RUNX3 | ko03070 | GG | -0.0017 | 0.0435 | Rectum |
| RUNX3 | ko03450 | AG | -0.0023 | 0.0368 | Rectum |
| RUNX3 | ko04141 | GG | -0.0014 | 0.0418 | Ileum |
| RUNX3 | ko04142 | AG | -0.0104 | 0.0251 | Rectum |
| RUNX3 | ko04974 | GG | -0.0032 | 0.0425 | Ileum |
| RUNX3 | ko05120 | AG | -0.0010 | 0.0397 | RC |
| RUNX3 | ko05120 | AG | -0.0010 | 0.0486 | Cecum |
| RUNX3 | ko05146 | GG | -0.0018 | 0.0135 | Ileum |
| RUNX3 | ko05150 | GG | 0.0015 | 0.0307 | LC |

**Supplementary Figure 1:** Receiver operating characteristic curve revealing substantial differentiation between AS cases and controls within the cohort (250 individuals).

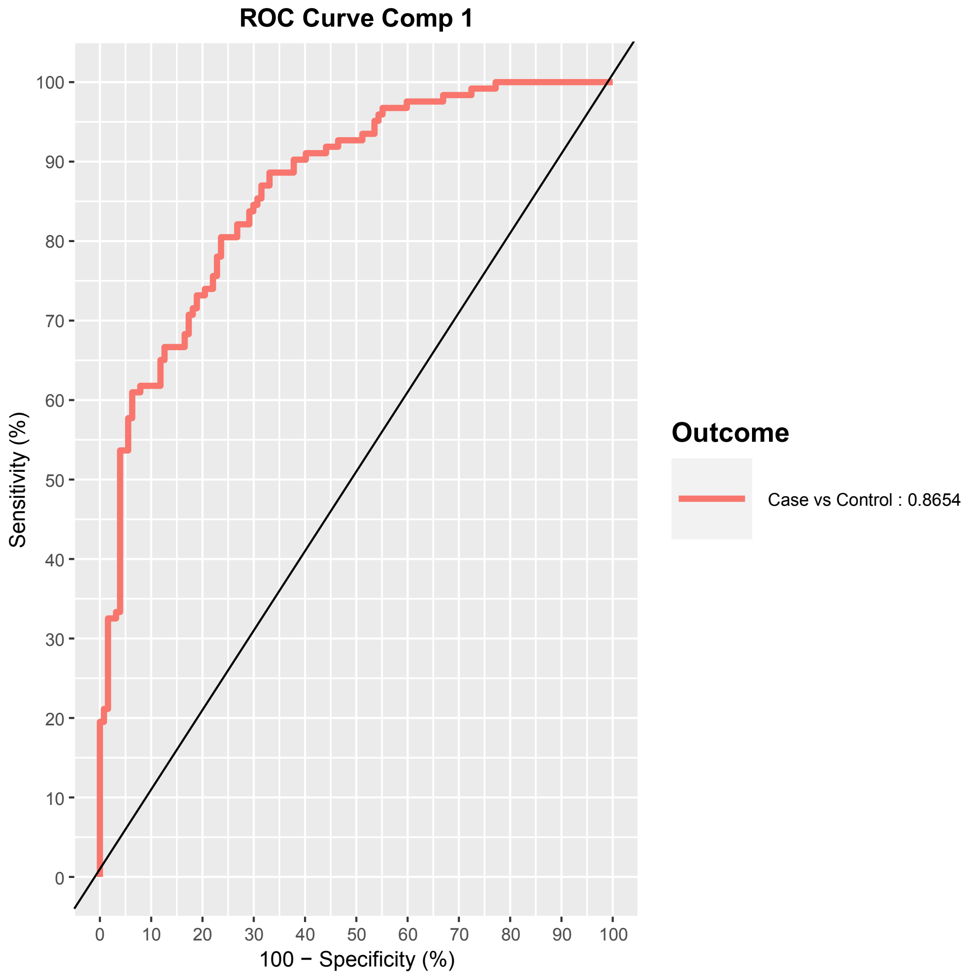

**Supplementary Figure 2:** Genetic-relatedness dendrograms of the strain population for each of the dysbiotic bacterial species in AS cases, as identified in Figure 1B. Samples without sufficient read coverage for these species were excluded. Each data point represents an indiviudal sample, which may consist of multiple strains. AS status was unable to distinctively separate the strains present in AS cases compared to healthy controls, indicating that gut dysbiosis may primarily be attributed to differential species abundance with no additional strain-level effects.

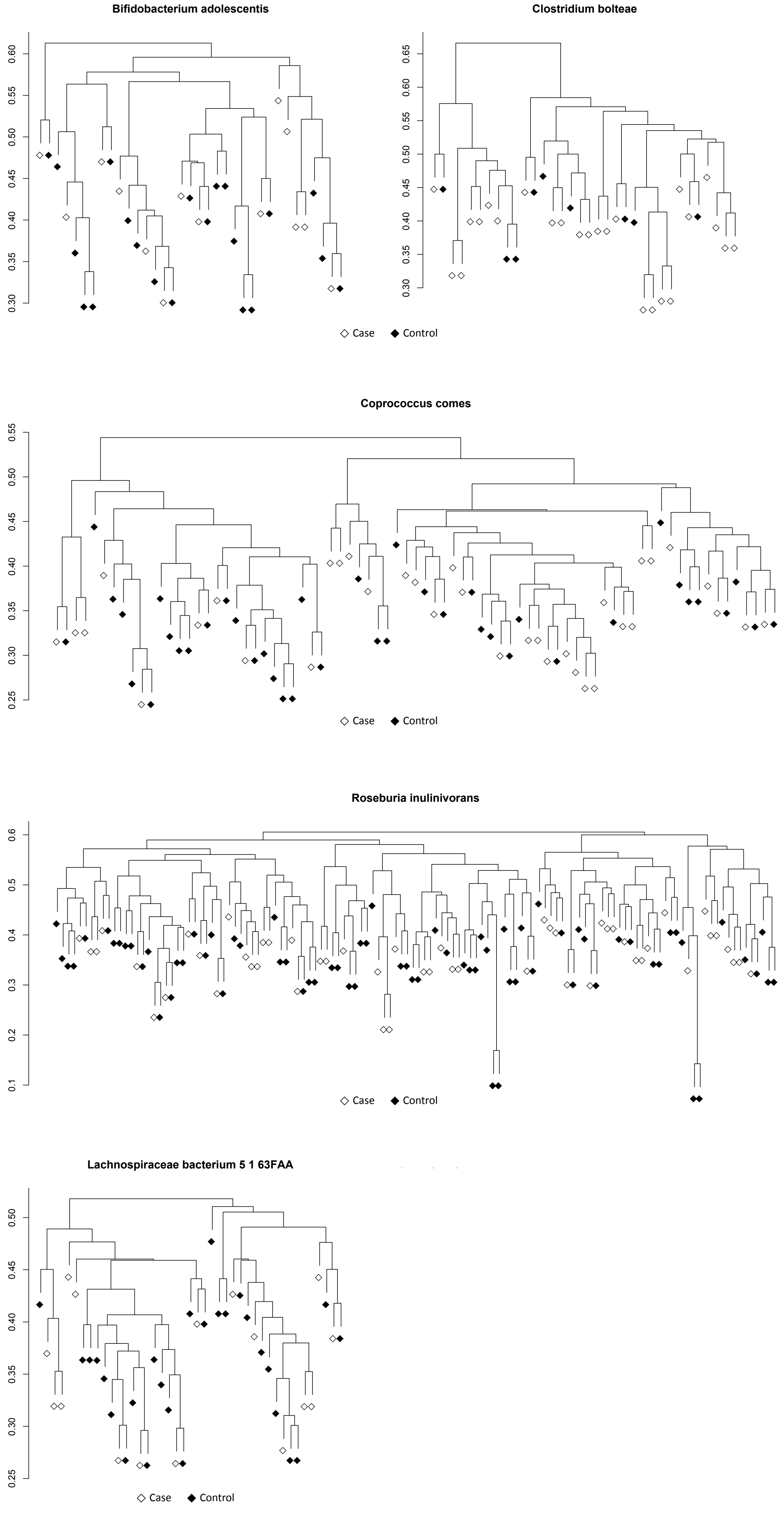

**Supplementary Figure 3:** sPLSDA and PERMANOVA results measuring the effect of sulfasalazine treatment upon the microbiome of AS cases, revealing a non-significant effect, potentially due to power or sample size constraints.

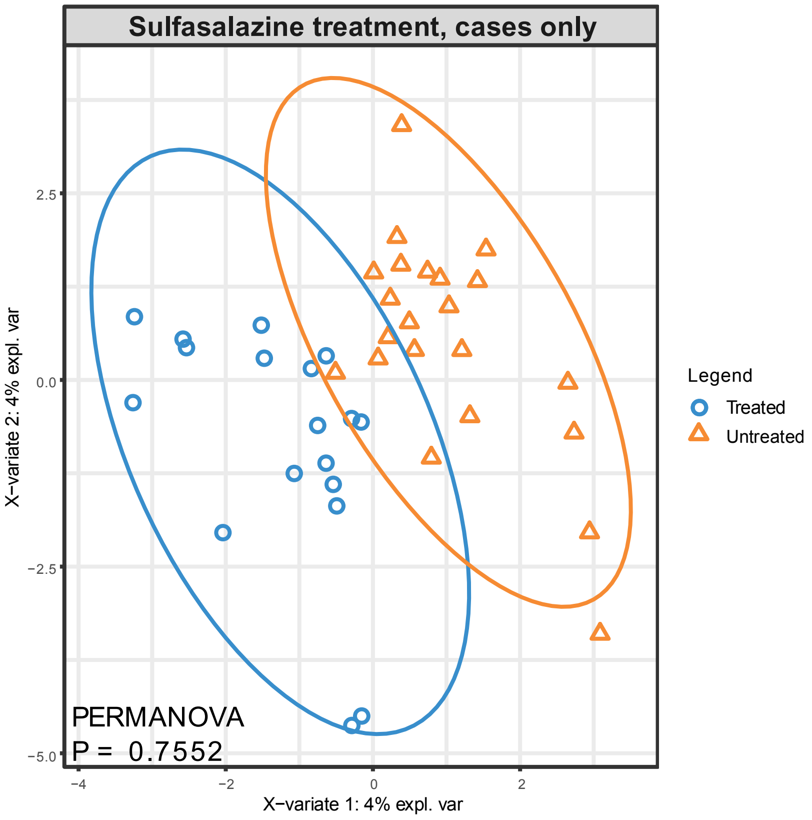

**Supplementary Figure 4:** Effect of TNFi therapy upon the dysbiotic **A.** bacterial species and **B.** KEGG Orthogroups which were commonly differentiated between cases vs controls in the discovery and validation cohorts, as shown in Figure 1. TNFi therapy appears to partially normalise dysbiosis, however statistical testing revealed non-significant differences between treated and untreated cases.

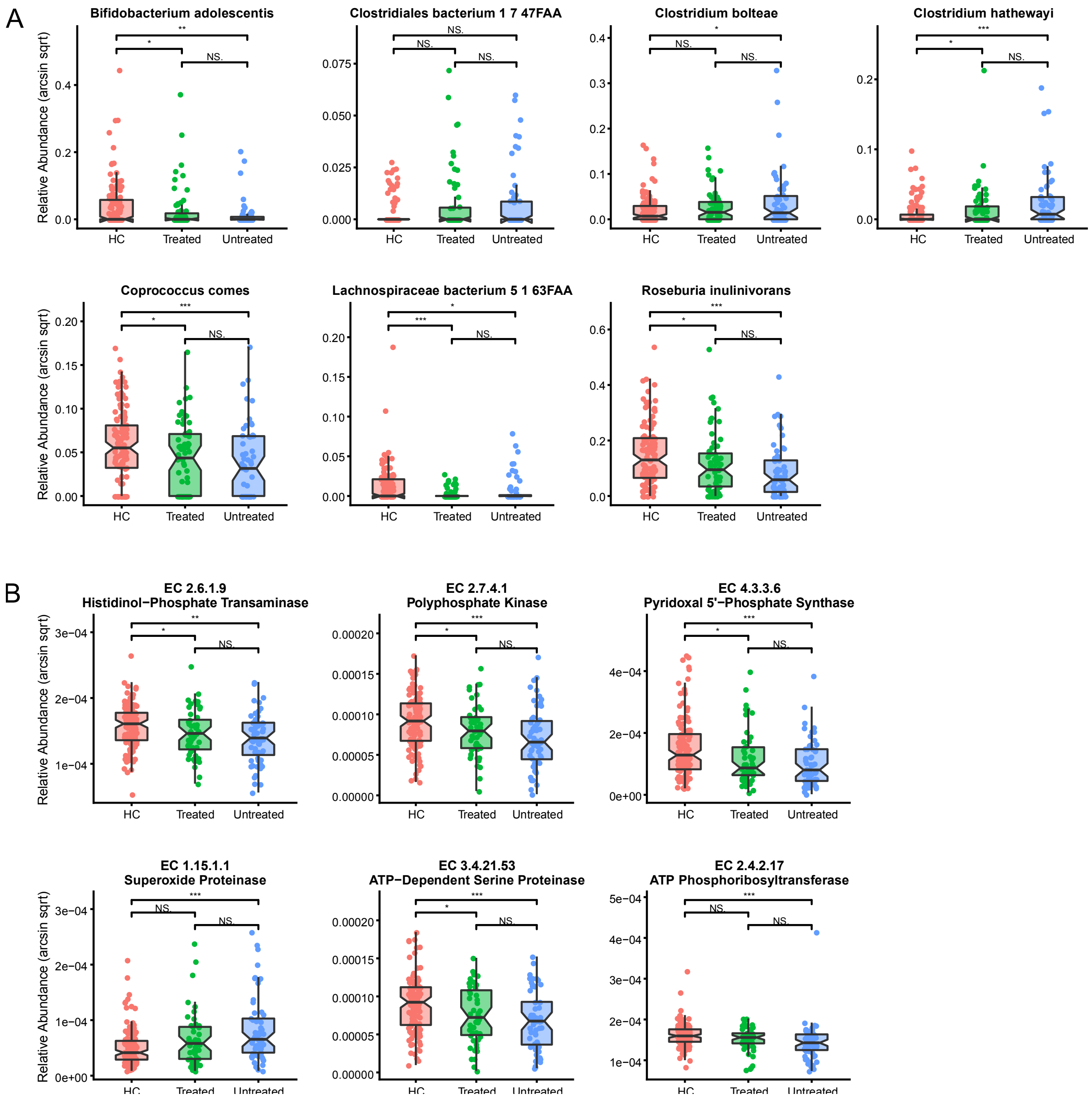

**Supplementary Figure 5:** Genetic-relatedness dendrograms of the strain population for each of the bacterial species modulated by TNFi therapy, as identified in Figure 2B. Samples without sufficient read coverage as well as species annotated as "unclassified" were excluded. Each data point represents an indiviudal sample, which may consist of multiple strains. No distinctive separation of strains present was noted, indicating that TNFi therapy may primarily affect species abundance with no additional strain-level effects.

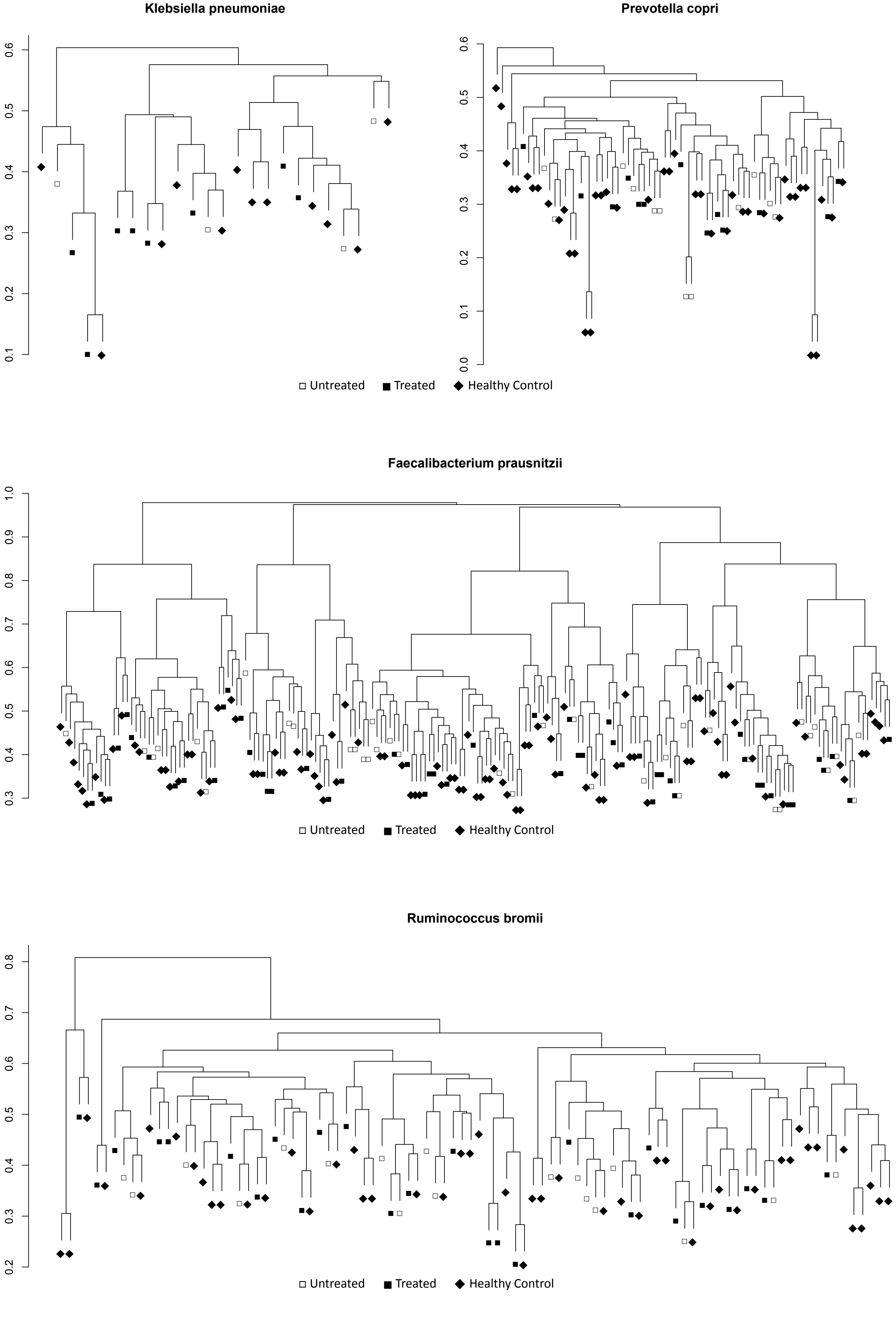

**Supplementary Figure 6:** Genetic-relatedness dendrograms of the strain population for differentially abundant bacterial species according to their RUNX3 genotype (rs11249215), as identified in Figure 3C. Samples without sufficient read coverage as well as species annotated as "unclassified" were excluded. Each data point represents an indiviudal sample, which may consist of multiple strains. No distinctive separation of strains present was noted, indicating that rs11249215 may primarily affect species abundance with no additional strain-level effects.

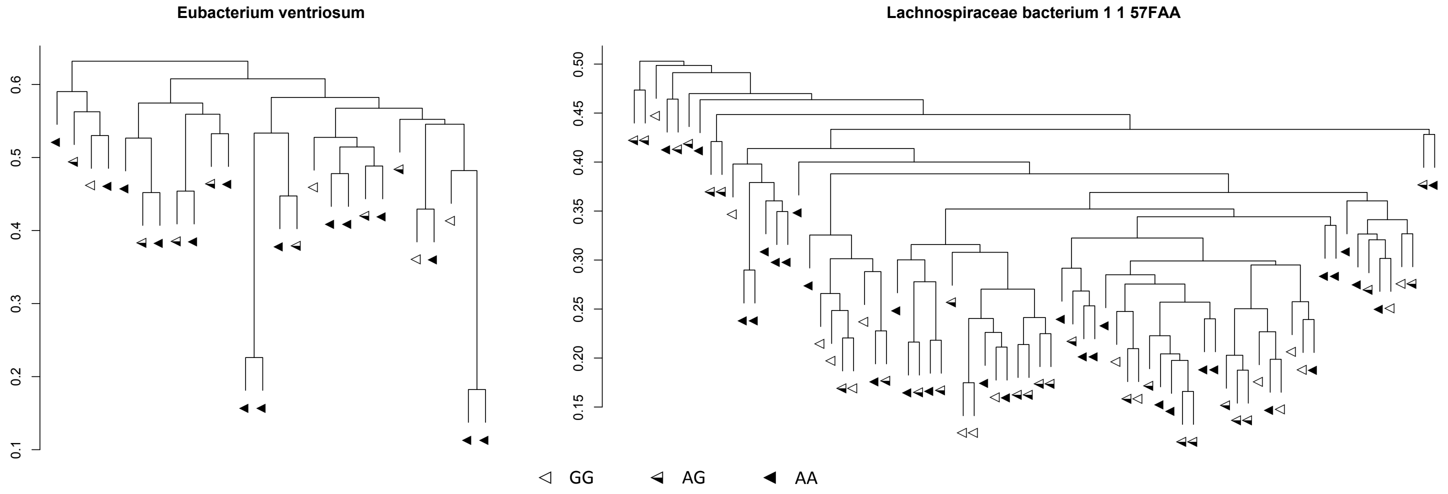
